# Metagenomics reveal allopatric speciation and higher connectivity among coastal vs. inland hypersaline lakes and solar salterns

**DOI:** 10.1101/2025.10.01.679725

**Authors:** Tomeu Viver, Juan F. Gago, Esteban Bustos-Caparros, Borja Aldeguer-Riquelme, Luis M. Rodriguez-R, Ana S. Ramírez, Luciana Albuquerque, Souad Amiour, Aharon Oren, Mehmet Burcin Mutlu, Stephanus N. Venter, Bonnie K. Baxter, María E. Llames, Bernardo González, Gustavo Rodríguez-Valdecantos, Horia L. Banciu, Matthew B. Stott, Fernando Santos, Brian P. Hedlund, Josefa Antón, Rudolf Amann, Konstantinos T. Konstantinidis, Ramon Rossello-Mora

## Abstract

Hypersaline environments, due to their discrete and geographically isolated nature, constitute ideal systems for studying evolutionary patterns and microbial diversification, and especially here when contrasting coastal with inland systems. Based on metagenomic comparisons of 25 hypersaline sites across 11 countries, we explored the influence of environmental factors, ionic composition, and geographic distance on their microbial community structures and taxa diversification. Our results revealed that microbial communities from coastal environments were taxonomically and functionally more similar to each other than to those from inland sites. A distance-decay relationship in the genetic relatedness, significantly more pronounced for the coastal sites, was observed among reconstructed metagenome-assembled genomes (MAGs), clearer at distances below 400 km, but still detectable across global scales up to 20,000 km. The 484 MAGs recovered, representing 284 distinct species, revealed a striking global ubiquity, with 62.5% of the species showed cosmopolitanism as were detected across multiple sites. The higher taxonomic and genetic similarity of coastal environments over the inland sites seems to reflect an environmental connection that may be related to the ocean current dynamics. Most cosmopolitan species showed clear allopatric differentiation, although few cases of a single globally dominant genomovar (average nucleotide identity, ANI > 99.8%) were also observed, especially for some *Haloquadratum* species. The findings suggest that coastal hypersaline systems are loosely constrained by geographic isolation, with clear signals of allopatric speciation at the mesoscale (tenths to hundreds of kilometers) that become blurrier at larger scales.

## INTRODUCTION

A longstanding challenge in microbial ecology is to elucidate the forces that drive the biogeography of microbial communities, and understand which selection drivers and processes support the dispersal and spatial assembly of communities [1,2]. Classical niche-assembly theory, as introduced by the Baas-Becking principle “everything is everywhere, but the environment selects,” proposes that microbial taxa have unlimited dispersal potential, with environmental conditions primarily determining local community composition [3]. In contrast, distance–decay patterns and insights from neutral theory highlight the importance of dispersal limitation, stochastic drift, and homogenization through dispersal, especially when gene flow is constrained by geographic or environmental gradients [4,5]. It is also important to note that microorganisms can evade some of these barriers by persisting in dormant viable states [6,7], allowing them to disperse via a wide array of vectors, including ocean currents [8], atmospheric aerosols such as clouds and dust [9,10,11], and animal hosts such as migratory birds [12,13], ultimately contributing to complex and often unpredictable biogeographic patterns. In this regard, the relative contribution of these dispersal mechanisms to the global distribution of halophilic microorganisms, particularly in discrete and geographically isolated hypersaline environments, remains largely unexplored.

To understand evolutionary dynamics within a single microbial species, dispersal must be viewed not only as a process of spatial redistribution but also as a key driver of gene flow, local adaptation, and lineage diversification [14]. The evolutionary consequences of dispersal depend on species-specific life histories, genome plasticity, ecological interactions, and the spatial configuration of their habitats. In microbes, where dispersal can be both frequent and global [15], evolutionary patterns are shaped by the combined effect between recurrent migration, genetic bottlenecks, and heterogeneous selection across environments [16]. When gene flow is restricted by geographic or ecological barriers, allopatric speciation can occur, promoting the divergence of lineages even within the same species [17].

In 2018, an international consortium of researchers formed the Halophile Sequencing Project (HSP) [18] with the aim to sample both coastal and inland hypersaline systems, such as lakes and solar salterns, and investigate microbial diversity, population maintenance, and evolutionary differentiation within these ecosystems. In contrast to the naturally occurring hypersaline lakes, solar salterns are semi-artificial, human-managed ecosystems designed for salt production through controlled cycles of seawater input, progressive salt concentration via evaporation, and salt precipitation. These cyclical processes generate highly selective, recurrent environmental conditions of increasing salt concentration to salt saturation, followed by dilution due to natural events (e.g., rain) or human activities, that strongly shape microbial communities. Every year, thousands of tons of seawater are pumped into coastal salterns and subsurface brines into inland salterns through human intervention, or may enter into natural coastal systems through sea spray or in natural inland hypersaline basins through streams dissolving evaporitic rocks serving as a continuous source of halophilic microorganisms. Previous research has demonstrated that salterns worldwide consistently harbour stable microbial communities [19,20], typically dominated by two primary lineages: the archaeal class *Halobacteria* and the bacterial family *Salinibacteraceae* within the class *Rhodothermia* [21,22,23,24]. It seems that the controlled and stable conditions of solar salterns promotes a reduced diversity but more robust community structures compared to naturally occurring hypersaline lakes [19,25]. Despite their low overall diversity, these hypersaline-adapted lineages exhibit considerable intraspecies richness. For instance, *Salinibacter ruber* may contain between 5,000 and 11,000 distinct genomovars (defined as clusters of genomes using ANI thresholds that typically fall between 99.2% and 99.8% [26,27]) within a single environment [26,28,29,30]. On the other hand, the most successful extremely halophilic species *Haloquadratum walsbyi* exhibits a remarkable worldwide homogeneity with very low genomovar diversity [31]. Hypersaline brine communities carry an important advantage for biogeography studies as their viability depends on high osmotic pressures (i.e., high salt concentrations) and seasonality (e.g., light intensity) that can be easily measured [32,33]. Most extreme halophiles cannot thrive in environments with salt concentration <15% [34], thus dispersal through air or water should be done in a metabolically inactive stage as dormant but viable cells [6]. Only the transport via animals, as colonizing hypersaline tissues like seabird nostrils with high salinity [12], would allow metabolically active cells to be dispersed. These facts make extreme halophiles both interesting and tractable for biogeography and allopatric speciation studies as evolution should take place in discrete habitats with minimal expecting evolution during dispersal.

In this study, we analysed a collection of 25 metagenomes from 19 hypersaline environments distributed with variable geographic distances across the globe. By integrating community-level metagenomics, which can reveal interspecies assemblage patterns, with the reconstruction of metagenome-assembled genomes (MAGs) to assess intraspecies diversity, we aimed to better understand the geographic distribution of microbial species and assess signs of allopatric adaptation. Additionally, we sought to identify the primary factors shaping microbial communities and driving speciation, focusing on environmental variables such as ionic composition, geographic distance, and potential dispersal mechanisms, including ocean current-mediated transport of halophilic microorganisms.

## MATERIAL AND METHODS

### Samples: origin and processing

In year 2018, an international research consortium formed the Halophile Sequencing Project (HSP), involving 19 researchers from 11 countries worldwide. The HSP team collected 25 samples from 11 countries (Table 1). For each sample, at least 1 L of brines was collected and used for the current metagenomic study and to isolate and describe new extreme halophilic species [18] and analyze the ionic composition. All samples were collected between September 2018 and January 2020, and were close to saturation levels for NaCl (25% to 37% total salts) at the time of sampling.

**Table 1.** Sampling location, date and salinity of the hypersaline sites in study. Salinity (%) given as measured using a Sper Scientific Salt Refractometer.

| Country | Location |  | Abbreviation | Hypersaline system | Sampling date | Coordinates | Salinity (%) |
| --- | --- | --- | --- | --- | --- | --- | --- |
| Spain | Mallorca | S'Avall | ES_AV | Coastal solar saltern | September 2018 | 39°19'28"N – 2°59'21"E | 36.2 |
|  |  | Es Trenc | ES_CM | Coastal solar saltern | September 2018 | 39°21'03"N – 3°00'45"E | 34 |
|  |  | Coco 1 – Mallorca | ES_CO1 | Coastal pool | May 2019 | 39°15'55"N – 3°03'05"E | 31 |
|  |  | Coco 2 – Mallorca | ES_CO2 | Coastal pool | May 2019 | 39°15'55"N – 3°03'05"E | 30 |
|  | Alicante | Santa Pola | ES_SP | Coastal solar saltern | May 2019 | 38°11'05"N – 00°35'08"W | 37 |
|  | Canary Islands | Janubio – Lanzarote | ES_LZ | Coastal solar saltern | June 2019 | 28°55'11"N – 13°49'15"W | 34.4 |
|  |  | Del Carmen – Fuerteventura | ES_FV | Coastal solar saltern | July 2019 | 28°21'58"N – 13°52'12"W | 35 |
|  |  | Arinaga 1 – Gran Canaria | ES_CA1 | Coastal solar saltern | October 2018 | 27°51'05"N – 15°24'07"W | 36 |
|  |  | Arinaga 2 – Gran Canaria | ES_CA2 | Coastal solar saltern | October 2018 | 27°51'05"N – 15°24'07"W | 34 |
| Algeria | Oum El Bouaghi | Sebkha Ezzemoul | DZ_AGR | Inland lake | January 2020 | 35°53'13"N – 6°30'50"E | 30 |
| Israel | Eilat | Eilat 3 | IL_EL3 | Coastal solar saltern | September 2018 | 29°33'29"N – 34°57'54"E | 33 |
|  |  | Eilat 6 | IL_EL6 | Coastal solar saltern | September 2018 | 29°33'29"N – 34°57'54"E | 34 |
| Portugal | Rio Maior | Rio Maior | PT_RMP | Inland lake | June 2019 | 39°19'34"N – 08°55'17"W | 32 |
| Romania | Transylvania | Fără Fund | RO_FF | Inland lake | April 2019 | 45°52'34"N – 24°04'03"E | 29 |
| Turkey | Konya | Tuz Lake 1 | TR_TUZ1 | Inland lake | June 2019 | 29°33'29"N – 34°57'54"E | 33 |
|  |  | Tuz Lake 2 | TR_TUZ2 | Inland lake | June 2019 | 29°33'29"N – 34°57'54"E | ND |
| USA | Utah | Great Salt Lake | US_CM | Inland saltern | July 2019 | 40°41'13"N – 112°22'33"W | 37 |
| Chile | Boyeruca | Lo Valdivia 1 | CL_LV1 | Coastal solar saltern | January 2019 | 34°41'56"S – 72°00'44"O | 26 |
|  |  | Lo Valdivia 2 | CL_LV2 | Coastal solar saltern | January 2019 | 34°41'56"S – 72°00'44"O | 26 |
| Argentina | La Pampa | Guatraché | AR_G | Inland lake | July 2019 | 37°42'59"S – 63°31'04"W | 33 |
|  |  | Colorada Chica | AR_CCH | Inland lake | August 2019 | 38°22'51"S – 63°25'41"O | 33 |
|  |  | Colorada Grande | AR_CG | Inland lake | August 2019 | 38°15'41"S – 63°45'02"O | 32 |
| New Zealand | Marlborough | Lake Grassmere | NZ_LG | Coastal lake | January 2019 | 41°44'33"S – 174°08'35"E | 37 |
| South Africa | Western Cape | Velddrif | ZA_VC | Coastal solar saltern | January 2019 | 32°44'03"S – 18°11'60"E | 34 |
|  | Bloemfontein | Soutpan | ZA_BS | Inland solar saltern | January 2019 | 28°45'21"S – 26°02'55"E | 25 |

### Ionic composition of the samples in study

To measure the ionic composition and concentration, samples were filtered through 0.22 μm hydrophilic PTFE filters, and the concentrations of fluoride (F^-^), chloride (Cl^-^), bromide (Br^-^), nitrate (NO_4_^-^), sulfate (SO_4_^2-^), phosphate (PO_4_^3-^), sodium (Na^+^), lithium (Li^+^), potassium (K^+^), ammonium (NH_4_^+^), magnesium (Mg^2+^) and calcium (Ca^2+^) were quantified by ion chromatography using a Metrohm, 850 ProfIC AnCat — MCS system, by the Technical Research Services of Alicante University (Spain). The salinities of the samples were measured using a Sper Scientific Salt Refractometer (model number 300006, Arizona, USA).

To investigate differences in ionic composition among samples, we conducted multivariate analyses in R (v4.2.3) using the vegan package [35] package. Similarity among samples based on ionic composition was assessed using Principal Component Analysis (PCA) on scaled data and Non-Metric Multidimensional Scaling (NMDS) based on Bray–Curtis dissimilarity. To evaluate statistically significant differences in ionic composition between sample groups, we applied permutational multivariate analysis of variance (PERMANOVA, adonis2) and analysis of similarities (ANOSIM, anosim), both performed on Bray–Curtis distance matrices.

### Metagenome analyses

DNA extraction from environmental samples was performed as detailed in Urdiain and colleagues [36]. Metagenomic samples were sequenced using an Illumina NextSeq500 instrument (2 x 150 bp, paired-end reads). Bbduk within the Bbtools package (http://bbtools.jgi.doe.gov/) was used to trim the paired-end reads with the following options: ktrim=r, k=28, mink=12, hdist=1, tbo=t, tpe=t, qtrim=rl, trimq=20 and minlength=100. Abundance-weighted average coverage and alpha-diversity indices of the metagenomes were estimated using Nonpareil v3.303 [37]. Beta-diversity analysis was determined using MASH distance v2.2.2 [38] and visualized in an NMDS plot using vegan v2.6 [35].

Trimmed reads were assembled with the SPAdes v3.13.1 [39] assembler using metaSPAdes mode and MEGAHIT v1.2.9 [49] with default parameters. Gene prediction from assembled contigs with >500 bps length was conducted using Prodigal v2.6.3 [41]. All predicted proteins from all metagenomes were compared using an all-versus-all DIAMOND blast tool v0.9.31 [42] and shared reciprocal best matches in all pairwise genome comparisons were identified using *rbm.rb* script [43] using a 50% sequence identity cut-off over 50% or more of the query sequence length. The orthologous group (OG) proteins were identified using *ogs.mcl.rb* script and the *Table.prefScore.R* script [43] was used to estimate the frequency of each OG in a given set of samples (coastal vs. inland samples). The representative sequence from each OG was annotated using the COG classifier v1.0.5 tool (https://github.com/moshi4/COGclassifier). Amino acid identity (AAI)-based similarity was visualized using NMDS plots generated with the vegan R package, applying a range of identity thresholds from 50% to 90%.

### MAG reconstruction and relative abundance

From the assembled files, contigs larger than 2,000 bps were binned using MaxBin v2.1.1 software [44] and MetaBAT v2 software [45]. Completeness and contamination values from MAGs were calculated using the CheckM2 tool v0.1.3 [46]. MAGs with completeness >50% and contamination <10% were selected for further analysis. dRep v3.4.0 was used to de-replicate the MAGs in species at 96% ANI value (using the FastANI algorithm [47]) and minimum level of overlap of 20% [48]. Representative MAG from each species was selected manually considering the higher completeness and lower contamination values. The phylogenetic placement of the representative MAGs was analysed using the GTDB-tk v2.4 tool with the GTDB Release R220 [49]. Species abundance was estimated using the representative selected MAG as a reference. For each metagenomic dataset, the MAG sequencing depth was estimated per position (Bowtie; bedtools) [50,51] and truncated to the central 80% values (i.e., removing the top and bottom 10% positions by depth) using the *BedGraph.tad.rb* script [43], a metric hereafter termed TAD80 (truncated average sequencing depth). Abundance was estimated as TAD80 normalized by the genome equivalents of the metagenomic dataset determined using the MicrobeCensus tool v1.1.0 [52]. A genomospecies was considered to be present in a sample if the TAD80 was non-zero and showing a sequencing breadth ≥ 20% (i.e., at least 20% of the genome covered by metagenomic reads). In order to estimate the frequency of a species in a given set of samples (i.e., accounting for the geographic distance or environmental characteristics) compared to the rest, we used the *Table.prefScore.R* script [43].

### Intra-species genome analysis

The average nucleotide identity (ANI) and the fraction of the genome shared between MAGs belonging to the same species were determined based on BLAST searches using *ani.rb* script [43]. Intra-species pangenome analysis was conducted following the methodology described by Conrad and colleagues [53]. Briefly, predicted genes were clustered into orthologous groups (OGs) at 90% nucleotide identity across at least 70% of the query sequence length using MMseqs2 v13.45111 [54], according to the guidelines provided in the pangenome workflow manual reported in https://github.com/rotheconrad/F100_Prok_Recombination. To assess gene abundance, metagenomic reads were mapped to the reference OG sequences using the BLASTn mode of Magic-BLAST+ v2.12.0 with default parameters [55]. Reads with a sequence identity >95% and an alignment covering >90% of the query read length were selected. For each gene, the resulting sequencing depth values were truncated to the middle 80% (TAD80) (i.e., the upper and lower 10% of outliers were removed) using a Python script from https://github.com/rotheconrad/00_in-situ_GeneCoverage [29]. For normalization, TAD80 values for each gene were divided by the mean TAD80 value across all OGs within each sample. To evaluate intra-species gene content variation across samples, pairwise dissimilarities based on normalized TAD80 values were calculated using the Bray–Curtis distance metric, implemented in the vegan R package. Additionally, Jaccard distances were computed based on gene presence/absence patterns.

## RESULTS

### Brines, ionic composition and geographic distance

The 25 brines were grouped into two categories according to their water source origin (Table 1 and Figure S1): **(i) Coastal (15 samples)**, in which brines originate from the direct concentration of seawater, including locations from the Spanish Mediterranean (S’Avall and Es Trenc from Mallorca, and Santa Pola from Alicante) and Atlantic littoral (Arinaga in Gran Canaria, Del Carmen in Fuenteventura, and Janubio in Lanzarote from the Canary Islands); from Israeli Aqaba gulf coast (Eilat); from the Pacific coast of Chile (Lo Valdivia); from the Atlantic coast of South Africa (Velddrif); and from the New Zealand South Pacific coast (Lake Grassmere). All were solar salterns with human-driven water feeding. In addition, we collected brines from ephemeral coastal ponds or “Cocos” (2 samples), which are small ponds adjacent to the seashore that are seasonally filled with seawater by sea spray; brines are formed due to evaporation in summer and get diluted due to rain in autumn and winter; both Coco samples were collected in the south coast of Mallorca. **(ii) Inland (9 samples)**, whose groundwater brines mostly originate from halite dissolution, and include samples from the solar salterns of Portugal (Rio Maior) and central South Africa (Bloemfontein), and the naturally occurring lakes in Romania (Fără Fund); the central Turkish region (Lake Tuz); the North African region of Algeria (Sebkha Ezzemoul); the North American Great Salt Lake (in Utah); and three Pampean lakes from Argentina (Colorada Chica, Colorada Grande and Guatraché).

All ionic compositions were consistent with a thalassohaline origin, being predominantly composed of sodium salts predominantly NaCl (Figure 1A and Table S1). Due to their seawater origin, coastal brines and Coco samples contained additionally a significant proportion of magnesium salts (MgSO_4_ and MgCl_2_). In contrast, magnesium salts were four to five times lower in the inland sites, mirroring the evaporitic rock composition source of the dissolved salts, with strong predominance of sodium salts (NaCl) and higher concentrations of calcium and lithium. No significant differences were found in phosphate concentrations between coastal and inland samples. Ordinations using NMDS and PCoA based on ionic composition (Figure 1B and Figure S2, respectively) revealed a clear separation between coastal and inland environments, supported by a significant ANOSIM result (R = 0.7, p = 0.001). PERMANOVA analysis indicated that 55% of the variation in ionic composition could be explained by the sample origin (Figure S2).

**Figure 1.**
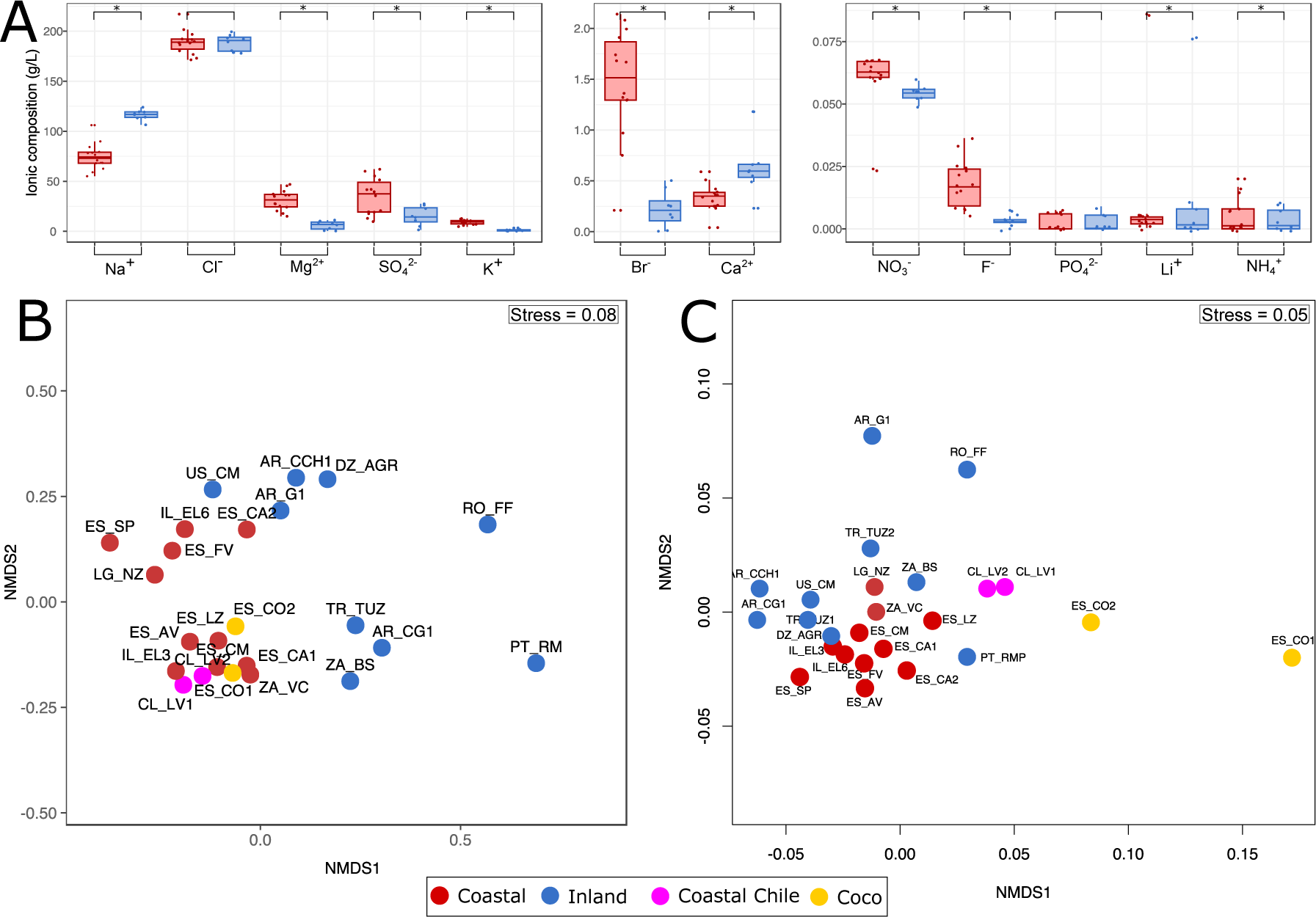
Similarity among all samples determined in this study. (A) Comparison of ionic composition (g/L) between coastal (red) and inland (blue) samples. (B) Non-metric Multidimensional Scaling (NMDS) based on ionic composition. (C) NMDS analysis of MASH-based distances (beta diversity index). Each point on the plot represents an individual sample, with red points indicating coastal and blue points indicating inland samples. An asterisk (*) marks statistically significant differences determined by the Wilcoxon test (p-value < 0.05).

Metagenome sizes after trimming ranged from 5.2 to 15 Gbp (average 11.5 ± 2.3), except the Santa Pola (ES_SP) dataset that was sequenced with 60 Gbp output (Table S2). As estimated by Nonpareil, all datasets showed a sequencing coverage ranging between 89 and 98%, and the variation observed in the sequence diversity index (*N_d_*) was dependent on sample (Figure S3A). The *N_d_* revealed that the coastal solar salterns exhibited lower diversity (17.6 ± 0.9) than the inland hypersaline lakes (18.2 ± 0.6; note that *N_d_* is in natural log scale), with statistically significant difference (Wilcoxon p-value < 0.05) (Figure S3B). Inland samples, such as those from Tuz Lake, reached *N_d_* values up to 18.9. The Coco samples also exhibited high *N_d_* values, with 18.1 and 17.4 in ES_CO1 and ES_CO2.

Beta diversity analysis based on MASH distance values (Table S3) revealed that coastal sites tend to share greater similarity among themselves when compared to inland sites, which displayed higher within-group variability. Betadisper analysis confirmed this pattern, with average distances to centroid of 0.053 for coastal and 0.075 for inland samples (Figure 1C). An ANOVA test indicated that this difference in dispersion was statistically significant (F = 7.96, p = 0.01). When assessing coastal and inland brines separately, we did not detect significant correlation between microbial community composition based on metagenomic MASH distance (Figure 1C) and ionic composition (dbRDA significance p-value > 0.7). In contrast to clustering based on ionic composition, the Chilean and Coco samples were exceptions as both sites clustered apart. Due to these differences, Chilean and Coco samples are treated separately below as their different community composition is related to their different functioning.

We detected a clear distance-decay of MASH overall relatedness among whole communities with geographic distance in kilometers (Figure 2). Both parameters exhibited significant correlation when analyzing the coastal sites independently (Procrustes correlation value of 0.73 and significance p-value < 0.01), but no significant correlation was observed for the inland samples (Procrustes correlation value of 0.57 and significance p-value 0.14) (Figure S4). Among the coastal salterns, a strong linear correlation with distance was evident for relatively short distances, up to 400 km (such as Arinaga and Eilat) reflected by an R^2^ value of 0.85 (Figure 2). The correlation decreased with increasing distances, being the lowest when considering the largest distances ranging from 5,000 to 20,000 km, but we still observed a similarity distance-decay pattern, with an R^2^ value of ∼0.2 (Figure 2). A decline in similarity with increasing geographic distance was also observed among the inland sites, but it was less pronounced. There, the R^2^ value dropped from 0.37 for distances ranging from 0 to 400 km to < 0.02 when including samples from distances of up to 20,000 km (Figure S5). Given the distinct distance decay patterns observed, samples were additionally grouped into three additional categories on the basis of their geographic proximity: from 0 - 400 km (small distance), 400 - 5,000 km (medium distance), and 5,000 - 20,000 km (large distance) (for details see Text S1 and Figure S1).

**Figure 2.**
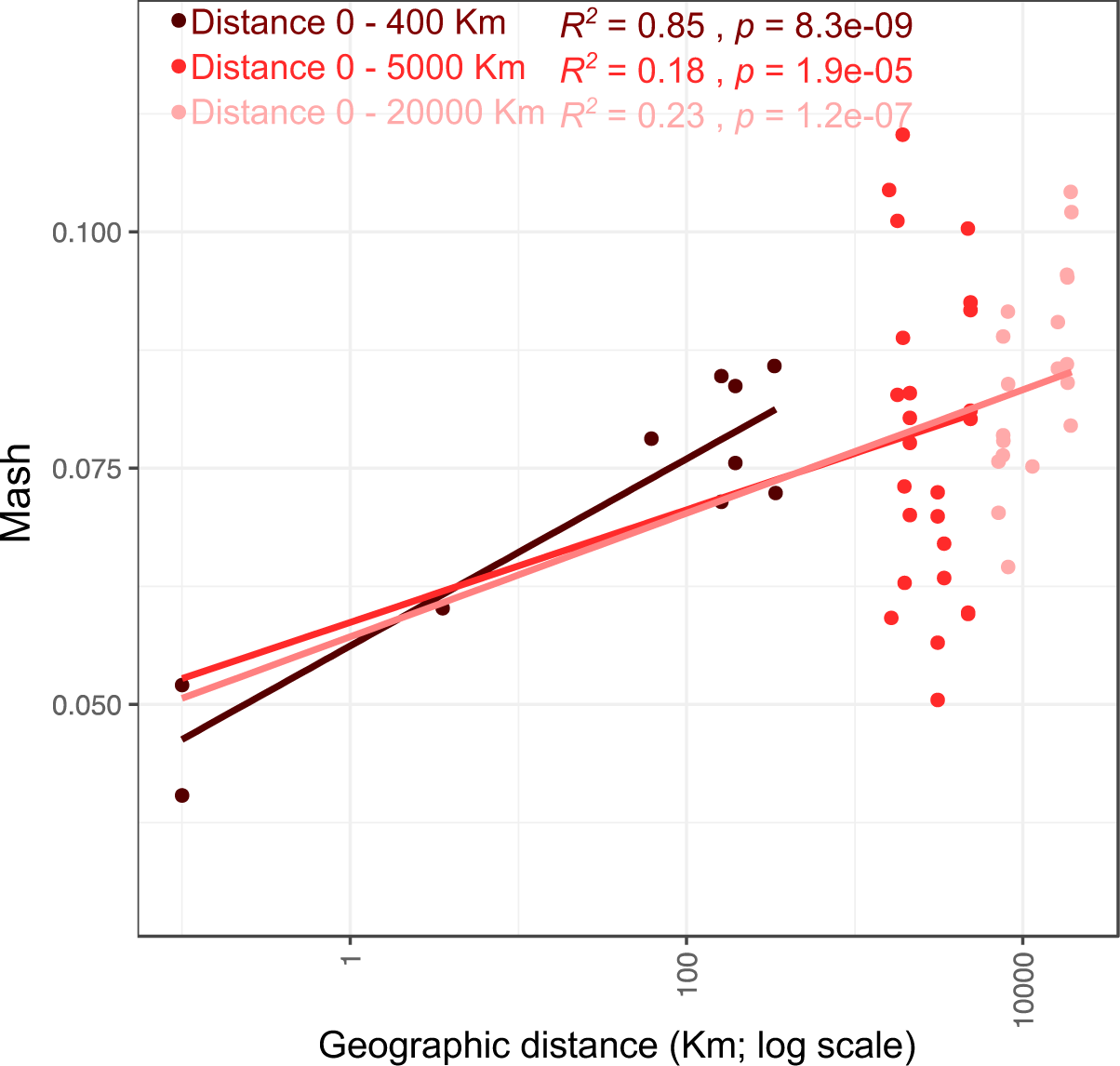
Distance-decay pattern of overall similarity among coastal microbial communities as a function of pairwise geographic distance (kilometers) based on the beta diversity index (MASH distance). Dark red represents all pairwise comparisons of samples within a 0–400 km range, while standard red extends the analysis to include additional samples up to 5,000 km, and light red further extends the analysis to distances of 20,000 km.

The average amino acid identity (AAI) among all samples from the assembled metagenomes, calculated with all predicted proteins with identities ranging from 50% to 90%, further confirmed that coastal sites shared higher level of similarity among themselves than with inland sites (Figure S6). Specifically, the median similarity among coastal sites was 90.8 ± 3.1%, whereas for the inland sites it was 83.4 ± 4.9%. From these predicted proteins, we identified 673,723 orthologous groups (OGs), and the comparison of their occurrence between coastal and inland environments revealed that 16.3% of OGs were more frequently found in coastal samples, 14.4% were more common in inland samples, and 69.2% showed no clear preference for either habitat (Figure S7A and B). Despite there being clear inland- or coastal-specific proteins, we did not find any special COG functional enrichment by type of saltern (inland vs coastal) nor we could detect any enrichment or selection of specific gene groups in any of the three protein categories (Figure S7C).

### Recovered MAGs and their taxonomic diversity and ubiquity

The 25 metagenomes rendered 484 good-quality MAGs, with an average estimated completeness of 76.25% (inter-quartile range [IQR]: 65.5%-86.4%), and an average estimated contamination of 2.43% (IQR: 0.45%-3.56%). Among them, 70 MAGs were of high-quality showing > 90% completeness (average 95 ± 2.8%) and < 7% contamination (average 1.5 ± 1.78%). After dereplicating the 484 MAGs at 96% ANI, we identified a total of 284 species (Table S4 and S5); based on GTDB-tk classification [49], 231 affiliated with the archaea and 53 with the bacteria. The most prevalent archaeal families were *Haloferacaceae* (n=97 species), *Haloarculaceae* (n=74), and *Nanosalinaceae* (n=49). Among these, 80.1% could be identified to the genus level, with *Halorubrum* (n=31), *Halovenus* (n=28)*, Nanosalina* (n=19), *Halobellus* (n=17) and *Natronomonas* (n=14) being the most species-rich (Table S5). Regarding bacterial species, 66% belonged to known genera and the most diverse were *Salinibacter* (n=11), *Roseovarius* (n=6), and *Spiribacter* (n=5). Moreover, members of unclassified higher taxa were identified within the classes *Bacteroidia* (n=7), *Bradymonadia* (n=3) and *Verrucomicrobiia* (n=2).

Metagenomic read recruitment using representative MAGs for the 284 species collection showed that the MAGs represented an average of 61.3% (IQR: 56.4%-74.4%) of the total metagenome, with the sample from Lanzarote (ES_LZ) recruiting the lowest (24.9%), and the sample from Algeria (DZ_AGR) the highest (87%) fractions of reads of their corresponding metagenomes (Table S5 and Figure S8). From the 284 species, 74 were locally restricted: 52 species were only detected in one sample, and 22 only in different ponds of the same site (i.e., in the two ponds from Eilat or Lo Valdivia, or the two Coco samples). In contrast, 210 species were detected in two or more distinct locations, with 61 of them showing high ubiquity, being present in 10 or more metagenomes. The most widely distributed species, in decreasing occurrence (Table S6), were *Halonotius pteroides* (23 sites), *Halorubrum* sp. Gsp_141_2 (22 sites), *Halonotius* sp. Gsp_89_2 (21 sites), *Sal. ruber* (20 sites), *Hqr. walsbyi* (18 sites), and *Haloquadratum* sp. Gsp_19_1 (17 sites). When evaluating the preference scores for the species between the coastal and inland sites (Figure 3A, 3B and Table S6), we observed a remarkable ubiquity for most species (62.5%). The analysis revealed that 78 species (29.2%) showed a clear preference for coastal sites, with 35 being exclusively coastal. Similarly, 74 species (27.7%) showed a preference for inland samples, and 65 were only detected at inland sites. A total of 115 species (43.1%) did not exhibit any preference for either environment. Among the genera with the highest species representation (e.g., *Halorubrum*, *Halobellus*, *Halovenus*, *Nanosalina*, *Natronomonas*, and *Salinibacter*), we identified both globally distributed species and species exhibiting clear preferences for either coastal or inland hypersaline environments (Figure 3C). We also observed a significant correlation between community structures and geographic distances using the Jaccard dissimilarity index among coastal samples (Procrustes correlation value = 0.67 and significance p-value < 0.01; Figure S9A), similar to that discussed above for whole-community relationships at the read level. This correlation was clearer in short distances up to 400 kilometers (R^2^ value = 0.64 (Figure S10)), but even extending to distant locations (5,000 to 20,000 km) the distance-decay still persisted with lower regression (R^2^ < 0.3). Such patterns were not observed for the inland sites (Procrustes correlation value of 0.38 and p-value > 0.05; Figure S9B and S11).

**Figure 3.**
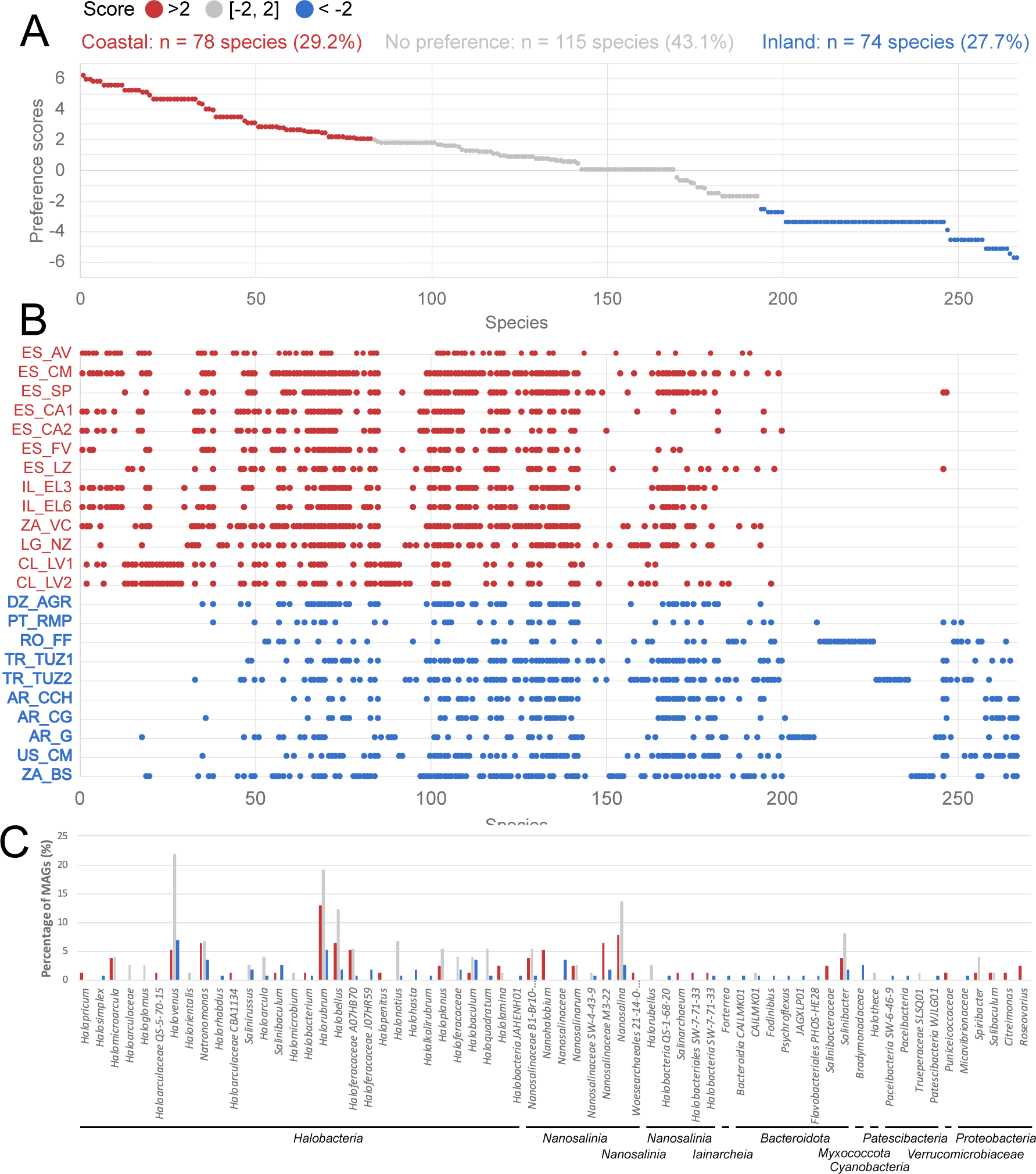
Preference scores of the recovered species in coastal and inland datasets. (A) Preference scores >2 indicate stronger preference for coastal samples (red) while scores <2 reflect preference for inland samples (blue). (B) Each point indicates the presence or absence of each species in the coastal (red) or inland (blue) samples included in this study and underlies the summary statistics shown in panel A. (C) Fraction of MAGs within each genus detected with preference for coastal (red bars), inland (blue bars), or no preference (grey bars).

### Intraspecific similarity patterns with geographic distance

Twenty-five species were binned from at least four metagenomes (Figure S12) and showed four distinct patterns of genome similarities based on ANI values (Figure 4 and S13):

**Figure 4.**
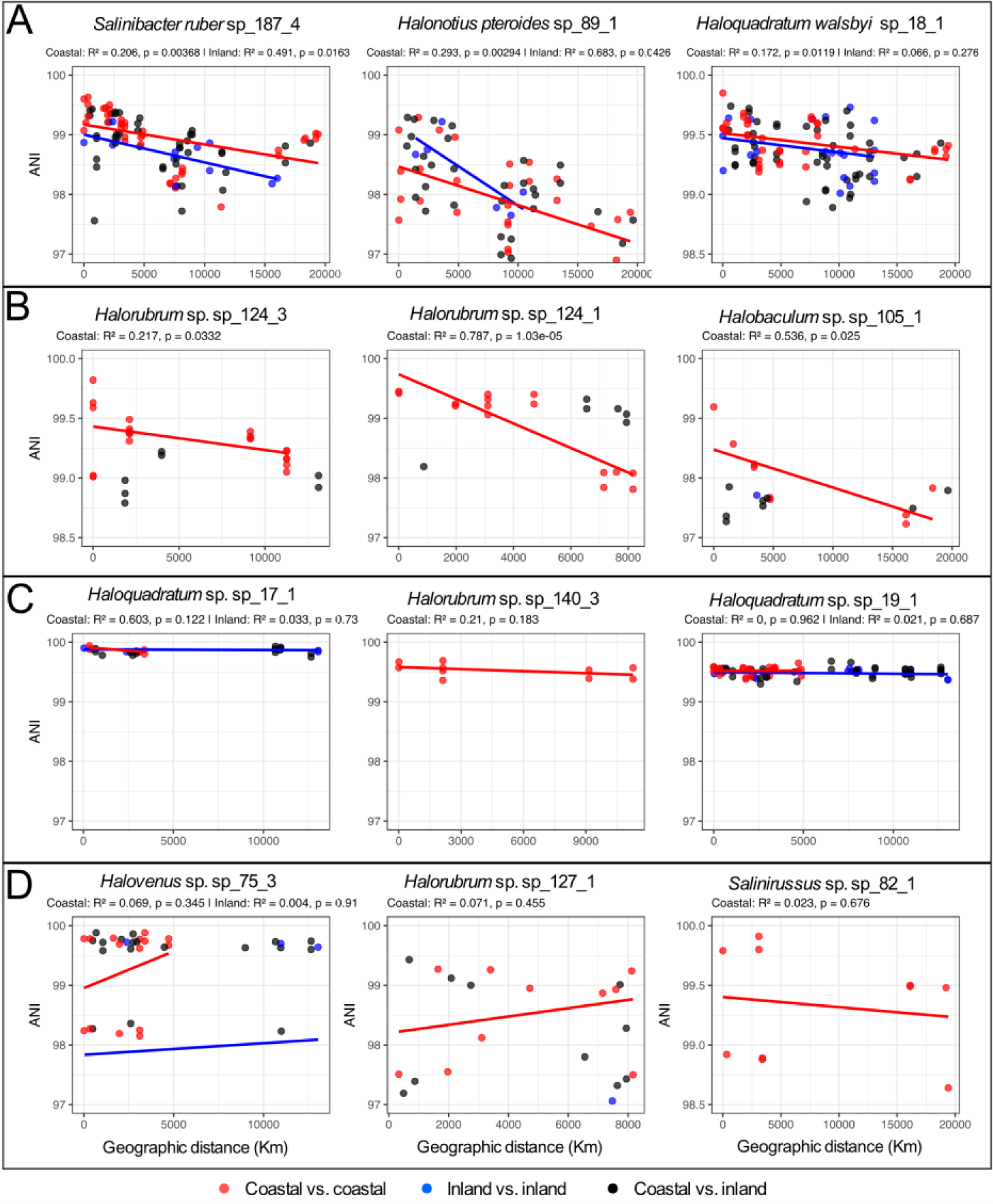
Relationships between intra-species average nucleotide identity (ANI) and geographic distances. (A) Species showing an ANI (y-axes) decrease with increasing geographic distance (x-axes). (B) Species showing the decrease only in coastal salterns, but with no correlation among inland sites. (C) Highly conserved species with ANI values >99. (D) Species with no correlation between ANI and geographic distance.

#### (i) Decreasing ANI with increasing distance

Some species, such as the widely distributed *Sal. ruber* (Gsp_187_4; 14 MAGs) and *Halonotius pteroides* (Gsp_89_1; 11 MAGs), showed a significant decrease in ANI with increasing geographic distance, regardless of their coastal or inland origin (p < 0.05). Alternatively, *Hqr. walsbyi* (Gsp_18_1; 15 MAGs) showed significant decrease in ANI only when comparing coastal sites (p < 0.05) (Figure 4A). Inland sites also showed a decrease but it was not significant (p = 0.276). For the three species, the lowest ANI values were observed when comparing the coastal and inland MAGs, with a maximum divergence of 96.9% ANI, and the patterns were consistent for both ANI values calculated using the whole MAG and only their orthologous core genes (Figure S13 and S14). The distribution of the identity values of the core genes across the three distance categories— short (0–400 km), intermediate (401–5000 km), and long (5001–20,000 km)—revealed a clear trend of decreasing ANI values with increasing geographic distance (Figure 5 and Figure S15). The results indicated stronger conservation of core gene sequences at local scales, contrasting with the reduction of their identities at larger distances. In addition, the variation in pangenome gene content, assessed using both relative abundance (Bray-Curtis index, Figure S16) and presence/absence (Jaccard index, Figure S17), showed a significant decline with distance, particularly between coastal samples (p-value < 0.05). Most of the gene content variation showing significant correlation either with distance or the inland vs coastal nature had unclassified functions, highlighting that a substantial proportion of the genes present in genomes of salt-adapted species remains poorly characterized. Among the functionally annotated genes, we did not observe a particular distribution of COG categories with the distance or nature of the samples (Figure S18). Although individual genes exhibited spatially structured abundance patterns (likely driven by environmental selection or dispersal constraints), the overall functional profiles did not display any relation on either geographic distance or environment (i.e., coastal vs. inland; Figure S19). These results suggested that while species-specific gene repertoires respond to spatial and ecological trends, these shifts did not result in consistent functional differentiation, but appeared to be functionally redundant, or the important functions remained hidden among the hypothetical and conserved hypothetical genes.

**Figure 5.**
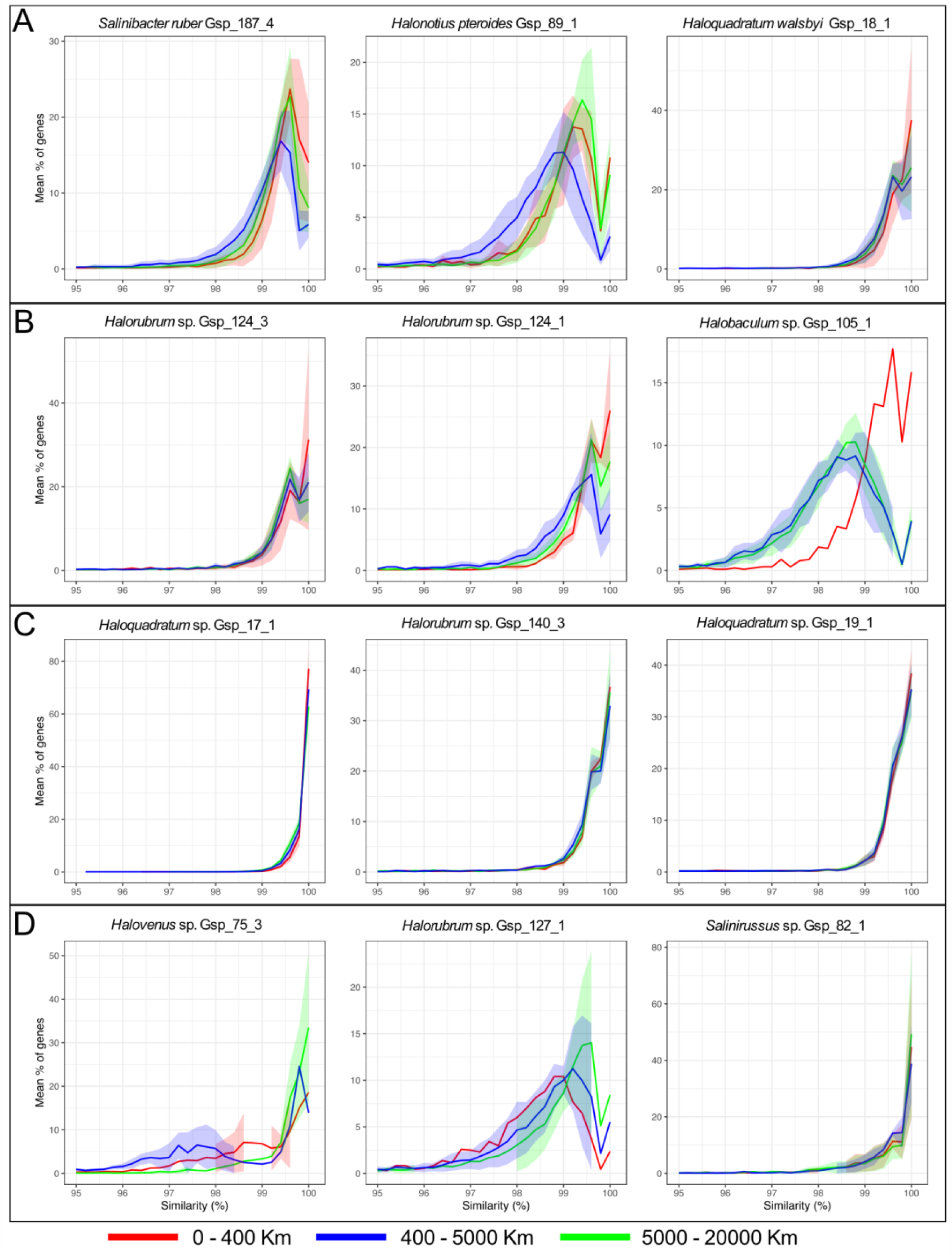
Intraspecies variation in core genes similarity across geographic distances. To assess the impact of geographic distance on genomic divergence within species, we compared the sequence similarity of shared core genes between MAGs recovered from different sampling sites. Pairwise comparisons were grouped by geographic distance into short-range (0–400 km; red), intermediate-range (401–5000 km; green), and long-range (5001–22,000 km; blue) categories. Each plot displays the average distribution of core gene similarity values for all pairwise comparisons within each distance group. Each line represents the distribution of core-gene similarity values binned into 0.2% intervals. The four panels are organized according to Figure 4: (A) species showing an ANI decrease with increasing geographic distance; (B) species showing the decrease only in coastal salterns, but with no correlation among inland sites; (C) highly conserved species with ANI values >99; (D) species with no correlation between ANI and geographic distance.

#### (ii) Increasing ANI distance with increasing distance only for coastal sites

Five additional archaeal species affiliated with the genera *Halorubrum* (Gsp_124_3, Gsp_124_1, and Gsp_139_1), *Halobaculum* (Gsp_105_1), and *Halobellus* (Gsp_157_1), which showed consistent patterns of genomic dissimilarity with geographic distance when comparing MAGs exclusively from coastal sites. These patterns were evident across multiple genomic analyses, including ANI (Figure 4B and S13B), core gene similarity (Figure 5B and S14B), pangenome structure (Figures S16B and S17B), and gene metabolic profiles (Figures S18B and S19B).

#### (iii) High ANI that remained constant with distance (no ANI variability)

We identified four species with high genome conservation with ANI values > 99.5% (that is, a single genomovar dominating) and two species with ANI values > 99.0% (potentially different, but closely related genomovars), in geographically distant locations or from both coastal and inland sites (Figure 4C and S13C). The most conserved was *Haloquadratum* Gsp_17_1, with an ANI value of 99.85 ± 0.05% in average and a minimum of 99.75%. This species was binned from various coastal western Mediterranean salterns, in Eilat and New Zealand, and also from inland lakes in Algeria, Turkey, and Argentina. Similarly, *Haloquadratum* Gsp_19_1, which was recovered from 12 different locations of both coastal (Canary Islands, Mallorca, Santa Pola, and Eilat) and inland areas (Algeria, Turkey, South Africa, and Argentina), exhibited an average ANI value of 99.5 ± 0.07% and a minimum of 99.3%. Supporting the highly conserved nature of the species, no significant differences were observed in the patterns of ANI distribution of the core genes in the different distance categories (Figure 5C). However, despite the high conservation, the MAGs from the different locations showed distinct gene content (i.e., were not clonal) and relative abundances in their pangenome (Figures S16C and S17C), but yet with no significant correlation with distance (p > 0.05), and no clear patterns of metabolic gene profiles (Figures S18C and S19C).

#### (iv) No correlation

We identified eight species with no correlation between ANI and distance (Figure 4D and S13D) and no correlation in the patterns of ANI distribution of the core genes in the different distance categories (Figure 5D) or the pangenome composition (Figure S16D and S17D). That is, for seven out of eight species in this category, we detected highly conserved MAGs (ANI > 99%) among geographically distant sites, but also MAGs with <99% ANI between close locations. For example, a group of *Halovenus* Gsp_75_3 MAGs from Es Trenc (Mallorca), Santa Pola, Eilat, Canary Islands, Turkey, Algeria, and Colorada Chica (Argentina) showed ANI values > 99.5%, but the conspecific MAGs form S’Avall (Mallorca) and Colorada Grande (Argentina) showed average ANI values of 98.2 ± 0.06% and 96.3 ± 0.11%, compared to the main group. Likewise, functional annotation of metabolic genes showed no enrichment or depletion of specific functional groups or COG categories (Figures S18D and S19D).

### Microbial diversity of the ephemeral brines of the Chilean salterns of Lo Valdivia and the Mallorca coastal hypersaline pools (Coco)

Similarly to the Cocos, the Chilean brines of Lo Valdivia can be considered as ephemeral as the area is flooded annually with seawater from the Pacific Ocean, which is subsequently evaporated in small evaporation and crystallizer ponds. The flooding returns the system to a nearly-initial situation every year, and therefore brines are not maintained stabilized as occurs in the majority of the solar salterns around the world. Due to this reason, both sites are considered apart. From the two Coco samples taken (ES_CO1 and ES_CO2; Table 1) biomass was similarly pigmented as other hypersaline sites (Figure S20). MASH distance (Figure 1C) and MAG diversity (Figure S21) showed ES_CO1 to be the most dissimilar from all samples, and ES_CO2 most similar to both Chilean samples. We binned 15 MAGs from ES_CO1 and 18 MAGs from ES_CO2, which were affiliated with genera commonly found in hypersaline environments (Table S5). Among all species detected from inland and coastal sites, 12 were exclusively from the Cocos, and 42 species were shared with other coastal and inland samples (Figure S22). Coco samples shared more species with coastal samples (14 ± 4 MAGs on average) than with inland samples (9 ± 7 MAGs on average; Figure S23B). The greatest overlap of Coco in species was observed with the two Chilean pools, CL_LV1 and CL_VL2, sharing 21 and 25 species, respectively (Figure S23A), and both sites also showed the highest similarity based on MASH distance (Figures 23C and 23D).

## DISCUSSION

As demonstrated in the past, salt concentration is a major deterministic factor shaping the community structures of hypersaline brines [30,56,57], consistent with high community similarities we observed. To challenge this established view and assess whether environments with broadly similar hypersaline conditions can still support intra- and interspecies variability, we sampled seawater-derived (thalassohaline) brines across wide geographical range. We have only considered environments with a direct or indirect/ancient seawater origin to avoid adding complexity to the study. Ponds in close proximity, such as the pairs taken within the salterns of Eilat and Arinaga, or between very close sites in Mallorca, Turkey, and Argentina, displayed the highest similarities in both metagenomic-inferred microbial community structures and gene composition as was expected. From a larger perspective, coastal samples, despite their global distributions, showed closer relationships compared to inland hypersaline sites with a similar global distribution. Communities inhabiting inland brines were then more diverse. The fact that communities inhabiting coastal sites were more coherent could be explained by either their similar ionic composition (e.g., higher magnesium) or to increased dispersal [6,7]. Mechanisms that could be responsible for global homogenization of microbes inhabiting coastal brines include transport in the atmosphere [9,10,11], via migratory birds [12,13], or currents. It seems that the conditions (over years) of osmotic variations below the threshold that would strongly disturb the community dynamics [58], act deterministically on how communities are structured, therefore responsible for sharing large amounts of taxa [59,60]. The coastal brines of Lo Valdivia and Cocos were exceptional given their highly divergent microbial compositions, and the most plausible explanation is their ephemeral hypersaline conditions, which are disrupted annually each cold/wet season, which could impede maturation of these communities, preventing their homogenization with other coastal sites.

Microbial communities inhabiting inland hypersaline environments showed higher dissimilarities, which could be attributed to more diverse brine composition or to geographic isolation, both of which could promote allopatric speciation. In turn, allopatric speciation would limit the efficiency of homologous recombination mechanisms [53], further limiting homogenization. In addition, as we speculated previously [61], hypersaline brines derived from groundwater dissolution of diapiric rocks could continually seed lakes with viable halophiles from the subsurface [62,63].

When taking geographic distance into account, we observed a clear similarity decay with distance in both genetic composition of the community and in the genomic identity of most cosmopolitan species. Again, this was clearer for coastal environments, yet communities in inland brines still showed some distance-decay, suggesting allopatric speciation. The distance-decay trend for the coastal environments was most evident at shorter distances (< 400 km) and became progressively weaker when comparing samples separated by 5,000 to 20,000 km, but these relationships were still more evident than those observed for inland ecosystems. This phenomenon can be explained by a combination of long-distance dispersal [64,65] and evolution. That is, the clear patterns at the mesoscale (from tens to hundreds of kilometers) can be explained by hydrological connectivity based on predictable currents carrying dormant halophiles over relatively small timescales. In contrast, the less clear distance-decay patterns at larger scales may reflect more complex ocean dynamics such as transport via downwelling, long-distance transport in deep currents, and upwelling, which may correlate less directly with distance and more with deep current residence time [66,67,68,69,70].

The significant redundancy in metabolic functions and species shared across all environments sampled evidenced that the osmotic pressure could be responsible as deterministic force [57]. For instance, ∼43% of the recovered species and ∼70% of the OGs showed no clear preference for their origin, suggesting a taxonomic and functional similarity in these hypersaline environments. As expected, the majority of these species affiliated with the same archaeal families, including *Haloferacaceae*, *Haloarculaceae*, and *Nanosalinaceae* being the most prevalent and the genera *Halorubrum*, *Halovenus*, *Nanosalina*, *Halobellus*, and *Natronomonas* dominating [23,71,72]. Additionally, the bacterial genera *Salinibacter*, *Roseovarius*, and *Spiribacter* were also prominent. As we demonstrated previously [24,26,31,58], thalassohaline hypersaline environments are commonly dominated by the same taxa with a large intra-taxonomic diversity that, due to the physiological versatility, guarantees their high prevalence [18,58], and explains why thalassohaline hypersaline sites share similar community structures globally.

The collection of MAGs encompassed 284 species with summed abundance in metagenomes ranging from 50% and 90% of the total microbial community, which is much higher than more diverse and variable marine and freshwater datasets [48]. From these species, 78 were exclusive to, or highly enriched in, coastal environments, whereas 74 were enriched in inland environments, indicating that many species (115) did not show any preference for coastal or inland environments. These patterns revealed that hypersaline environments worldwide not only exhibited phylogenetic and functional similarity at higher taxonomic levels [23], but also at the genus and species levels. For instance, *Salinibacter* and *Halorubrum* exemplify globally distributed generalists that co-occur with locally adapted, and often different species of the genus, species that are restricted to specific salterns [18,73]. Furthermore, intraspecific microdiversity studies suggested that generalist species often comprise multiple ecotypes with distinct functional capacities, enabling persistence across spatial and temporal salinity gradients [25,58].

The major outliers in the study were brines from the ephemeral hypersaline environments of Lo Valdivia on the Chilean coast and the two Cocos from the coast of Mallorca. The important taxonomic and genetic dissimilarities of both systems and the rest of samples may rely on the immature nature of these brines, which are yearly assembled from seawater carrying the presumably dormant extreme halophiles. It seems that when *de novo* assembling from seawater there is a period of community structure transitions, with unstable intermediate stages in their composition, that need prolonged periods of months of osmotic stability to reach the commonly found community structures of the coastal brines (Gonzalez-Mendez, unpublished results). The yearly disruption of the brines due to dilution with stream freshwater (in Lo Valdivia) or rainfall seem to occur before a robust mature community is be established. Consistent with the latter interpretation, we did not find any trace of the most abundant member of nearly all examined brines, *Hqr. walsbyi*, a fact that is also observed in the brines of the salterns of Cahuil [74] around 40 km north of Lo Valdivia, and probably being managed with a similar seawater feeding mechanism to that of the ephemeral Coco sites. It would be interesting to experimentally determine whether long-term stabilization of these Coco brines would result in communities that mirror those in stable hypersaline environments.

When focusing on the geographical patterns of the cosmopolitan species, distance-decay was evident for the most abundant and ecologically successful species such as *Hqr. walsbyi*, *Sal. ruber*, and *Halonotius pteroides*, in which the number of shared genes with high nucleotide identity (≥99.6%) from nearby sites (0–400 km) progressively decreased in both number and identity at intermediate and especially long-range distances. This pattern was also evident in both core and pangenomes, suggesting a clear genomic divergence with an apparent metabolic similarity, but probably with distinct ecological fitness.

While it seems that ocean dynamics might contribute to the homogenization of microbial populations across coastal sites, the observed genomic patterns also highlighted the role of allopatric speciation, suggesting that both dispersal-driven homogenization and geographic isolation may shape evolutionary trajectories at the intraspecific level. One of the additional points of interest in this study was the detection of a reduced number of species characterized by extremely high genomic relatedness even among geographically distant locations, as the highly clonal species of *Haloquadratum (Hqr.* GSP_17_1 and *Hqr.* GSP_19_1; not *Hqr. walsbyi*) that shared ANI > 99.8% worldwide. Although we cannot yet explain this phenomenon, it suggests that specific genomovars of some species may dominate worldwide and would probably be the best-adapted lineages in these systems [31] persisting in the environment for long periods and outcompeting other genomovars with lower ecological fitness. Another plausible explanation would be the especially effective worldwide dispersion of the same establishing genomovars given their broad ecological niche. However, this explanation would also imply that the clear allopatric speciation for the majority of the cosmopolitan species should be blurred. Future studies taking into account the trajectories and residence times of ocean currents together with larger number of sites will enlighten the role of oceans in homogenizing extreme halophilic populations.

## Data availability

The genomes and metagenomes generated in this study were deposited under the European Nucleotide Archive (ENA) accession code PRJEB45291 (the individual metagenome accession and MAG code has been listed in Table S2 and S7, respectively).

## Acknowledgements

Authors especially thank the Salinas de S’Avall, the Salinas d’Es Trenc and Gusto Mundial Balearides, S.L. (Flor de Sal d’Es Trenc), the Salt of the Earth Ltd., Eilat and the Interuniversity Institute for Marine Sciences of Eilat for (both by connection with A.O), the Dominion Salt in Lake Grassmere (by connection with M.B.S.), the State of Utah (by connection with B.H. and B.K.B), the local security authorities in the Oum El Bouaghi-Sebkha Ezzemoul (by connection with A.S.), the Ocna Sibiului city hall, Fără Fund Lake (in connection to H.L.B.) for supporting the access to the samples and facilitating the experimentation. HLB acknowledges the support of Dr. A. Cristea and Dr. Zs. Keresztes during sampling and sample preparation.

## Sampling permits

Brines collected from the Spanish Solar Salterns were authorized by the Spanish Ministry of Ecological Transition with the permit numbers: ESNC22 and ESNC27. Samples from Great Salt Lake were collected in accordance with permission from the State of Utah. Landowner permissions were obtained for sampling of the South African salterns. Local security authorities allowed sampling at the Oum El Bouaghi-Sebkha Ezzemoul hypersaline system in Algeria. Samples from Turkey were collected in accordance with the permission from the salt production site operators of Tuz Lake. For the samples collected in Argentina, the corresponding permit was issued by the “*Dirección de Recursos Naturales*” of La Pampa (Argentina), operating under the Ministry of Production (“*Subsecretaría de Asuntos Agrarios*”). Sample from Fără Fund Lake (Romania) was collected with the permission from Ocna Sibiului city hall. All sampling campaigns were done in accordance with compliance with the permit requirements in each country and/or region.

