## Supplemental Tables for "Metagenomics reveal allopatric speciation and higher connectivity among coastal vs. inland hypersaline lakes and solar salterns"

**Supplementary text S1:** Given the distinct distance decay patterns observed samples were additionally grouped into three in distance categories on the basis of their geographic proximity: from 0 - 400 Km, 400 - 5,000 Km and 5000 - 20,000 Km: **(i) small distance (0 - 400 Km)**: accounted for samples separated by <1 km (close ponds within the same saltern of Eilat and Arinaga; and two sampling points of Lake Tuz); separated by ~4 km as the two salterns in Mallorca (S'Avall and Campos); 61 to 195 km apart as the three Pampean lakes in Argentina (Colorada Grande, Colorada Chica, and Guatraché), as well as the three salterns in the Canary Islands (Arinaga, Del Carmen, and Janubio); and finally the Mediterranean salterns Santa Pola (Alicante) and Mallorca distanced by ~340 km. **(ii) medium distance (400 - 5,000 km)**: accounted pairs of samples from the Mediterranean, Canary Islands, Portugal, Algeria, Romania, Tuz Lake, and Eilat, as well as between the two South African samples. **(iii) large distance, (5,000 - 20,000 km)**, accounted for the geographical continental region pairs from the subcontinent's samples Europe, USA, South Africa, South America, and New Zealand. The largest distance was between Rio Maior (Portugal) and Lake Grassmere (New Zealand) with ~19,641 km (Figure S1 and Table S8 for detailed linear geographic distances between location pairs).

**Figure S1.** Geographical distribution of the hypersaline samples collected in this study. All samples were collected between September 2018 and August 2019. Coastal solar salterns or inland hypersaline lakes are marked with red or blue star, respectively. The distances given refer to Mallorca as a reference (geographic distances between pair of location have been reported in Table S8). Background colors indicate the geographic distance between samples, classified as short (0–400 km; green), intermediate (400–5,000 km; red), and long distances (5,000–20,000 km; blue).

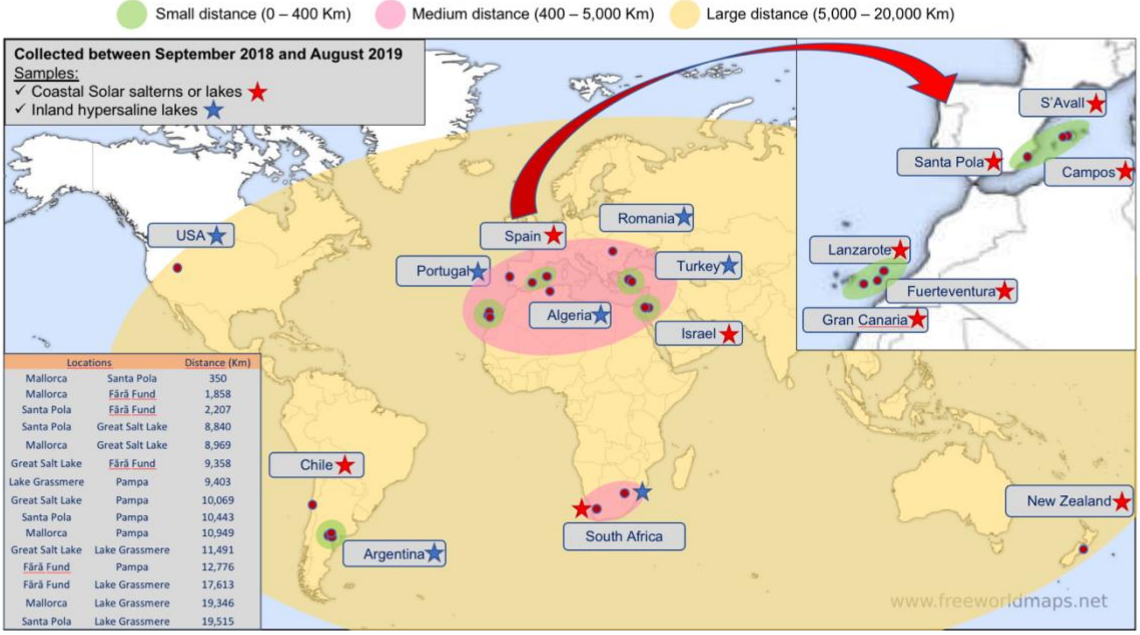

**Figure S2.** Similarity of ionic composition among all samples included in the study. (A) Non-metric Multidimensional Scaling (NMDS) analysis based on ionic composition. (B) Principal Component Analysis (PCA) based on ionic composition.

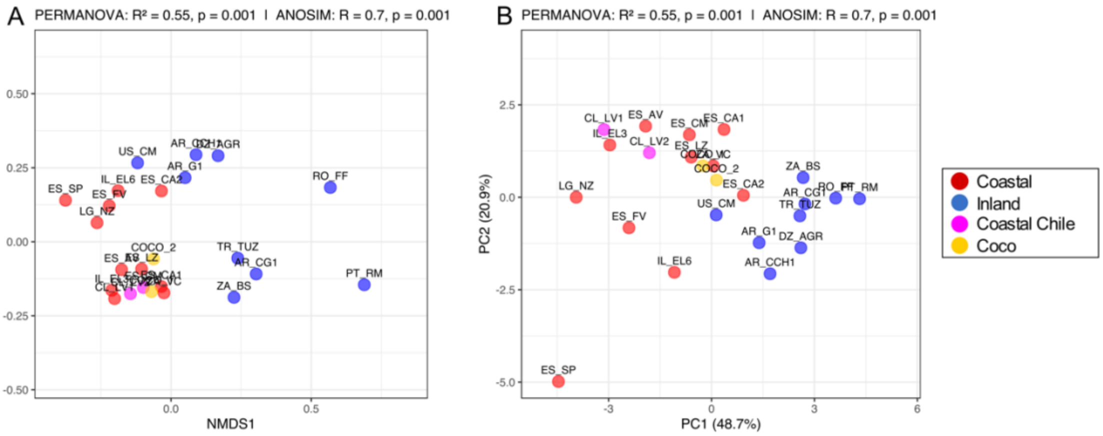

**Figure S3.** Metagenomic alpha diversity index based on Nonpareil ($N_d$) (A) for each individual sample, and (B) boxplot differentiated by coastal solar saltern and inland hypersaline lake origin.

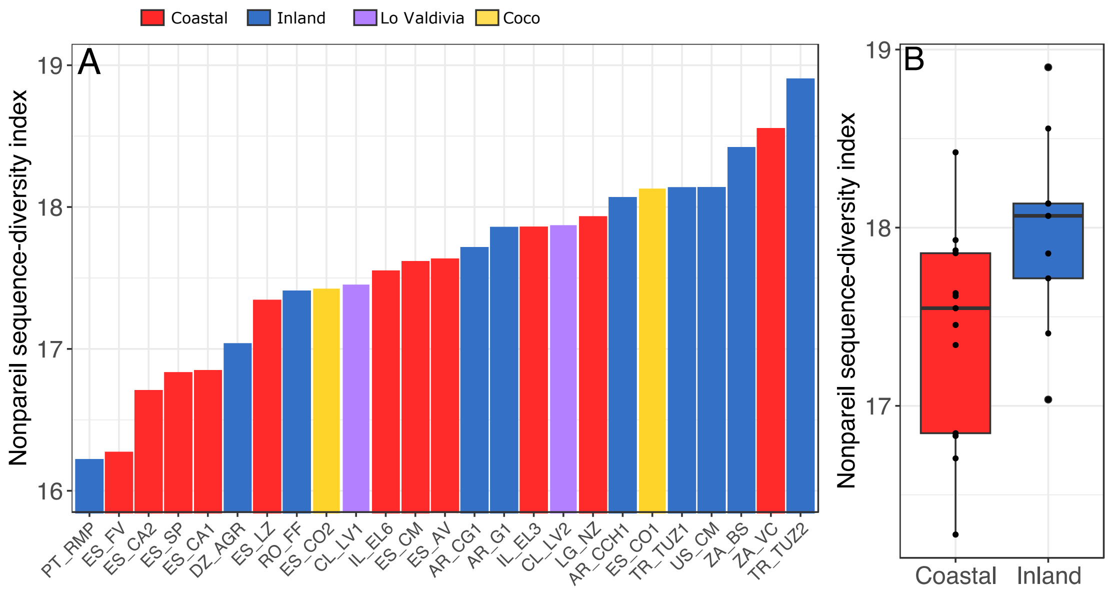

**Figure S4.** Procrustes statistical analysis comparing microbial composition through MASH distance values and geographic distance in kilometers of the coastal (A) and inland (B) samples independently.

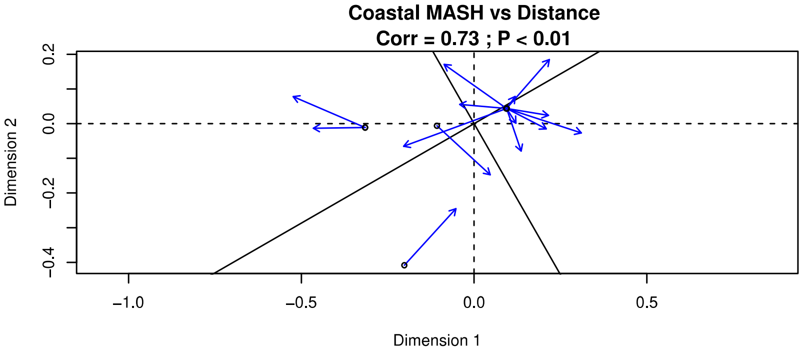


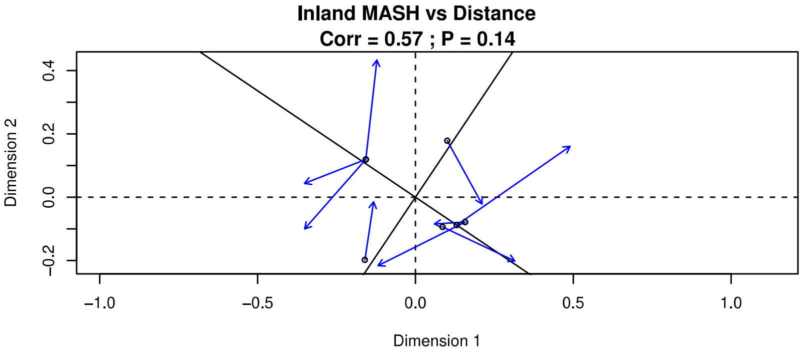

**Figure S5. Similarity among all inland hypersaline lakes based on MASH distance values in relation to pairwise geographic distance (kilometers):** Dark blue represents all pairwise comparisons of samples within a 0–400 km range, while standard blue extends the analysis to include additional samples up to 5,000 km, and light blue further extends the analysis to distances of 20,000 km.

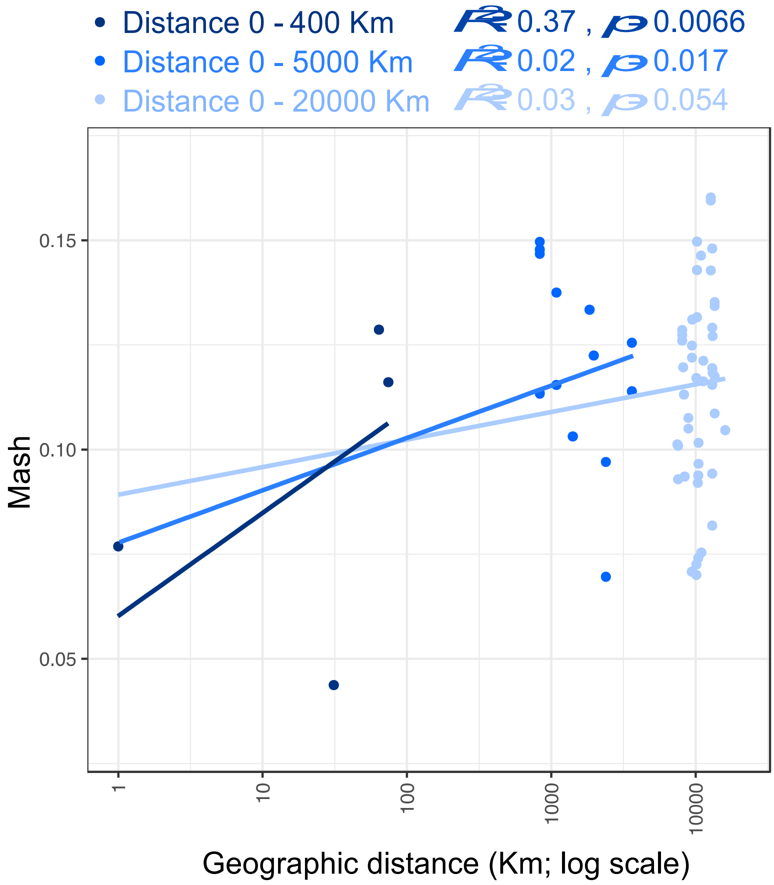

**Figure S6.** Average amino acid identity (AAI) among all samples considering different similarity thresholds, from 50% to 90%. Similarity was plotted with a Non-metric Multidimensional Scaling (NMDS) analysis.

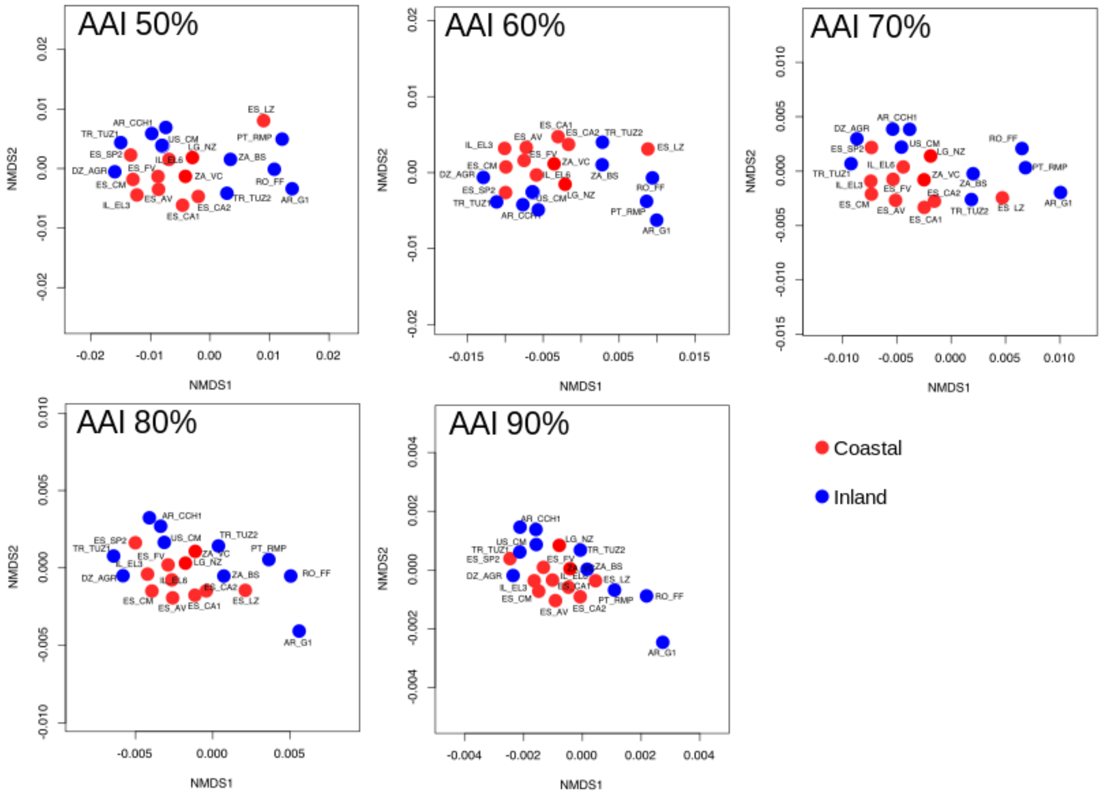

**Figure S7.** Preference scores of the protein orthologous groups (OGs) predicted from the assembled metagenomes. (A) OGs with preference scores >9 indicate a higher prevalence in coastal salterns (red), while scores <9 indicate a greater prevalence in inland samples (blue). (B) Each point represents the presence or absence of OGs across the samples included in the study. (C) Metabolic classification of the OGs based on COG functional categories (see legend), differentiating between coastal (red bars), inland (blue bars), or no specific environmental preference (grey bars).

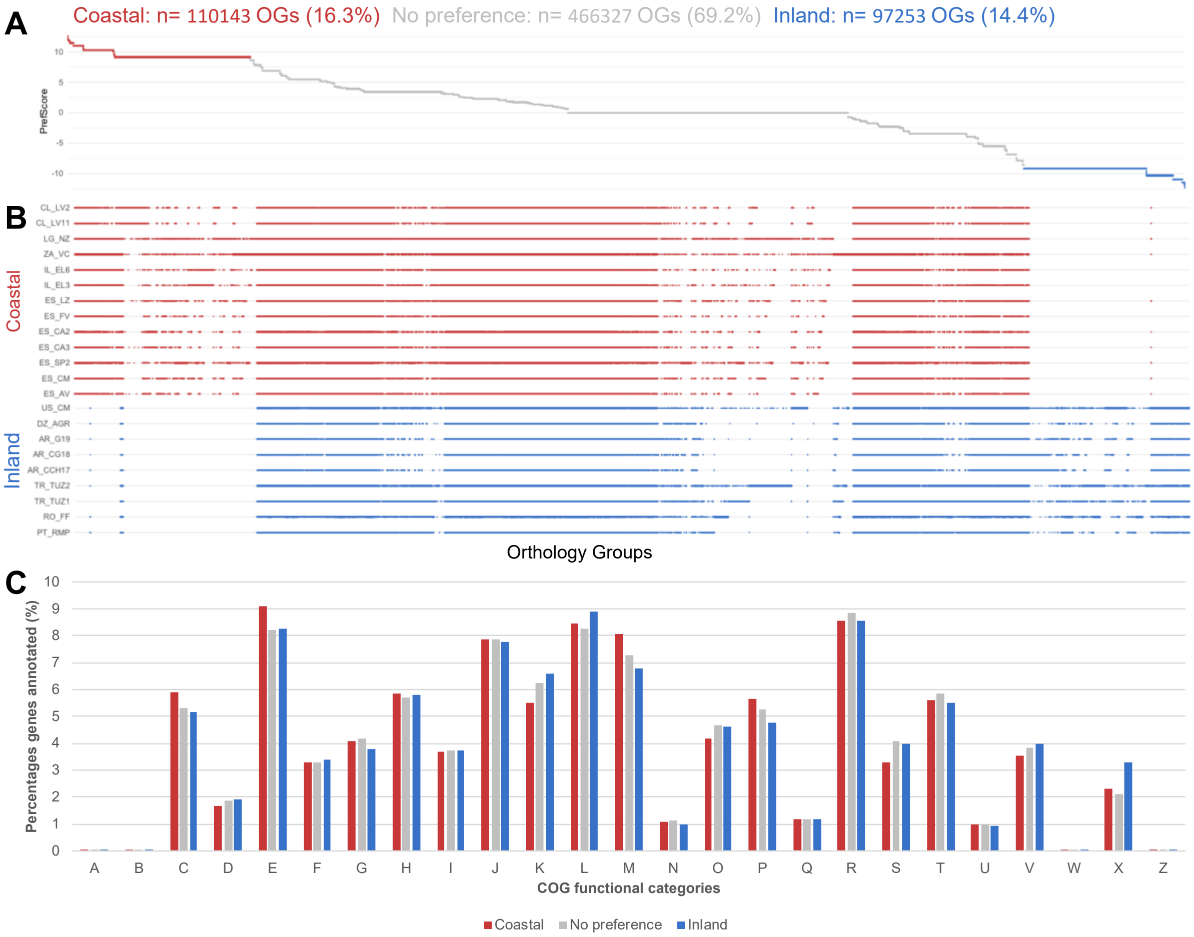


A RNA processing and modification
B Chromatin structure and dynamics
C Energy production and conversion
D Cell cycle control, cell division, chromosome partitioning
E Amino acid transport and metabolism
F Nucleotide transport and metabolism
G Carbohydrate transport and metabolism
H Coenzyme transport and metabolism
I Lipid transport and metabolism
J Translation, ribosomal structure and biogenesis
K Transcription
L Replication, recombination and repair
M Cell wall/membrane/envelope biogenesis
N Cell motility
O Posttranslational modification, protein turnover, chaperones
P Inorganic ion transport and metabolism
Q Secondary metabolites biosynthesis, transport and catabolism
R General function prediction only
S Function unknown
T Signal transduction mechanisms
U Intracellular trafficking, secretion, and vesicular transport
V Defense mechanisms
W Extracellular structures
X Mobilome: prophages, transposons
Z Cytoskeleton

**Figure S8.** Total abundance of species detected by MAGs in each metagenome. Bars are color-coded by sample origin: red for coastal, blue for inland, purple for Lo Valdivia, and yellow for Cocó samples.

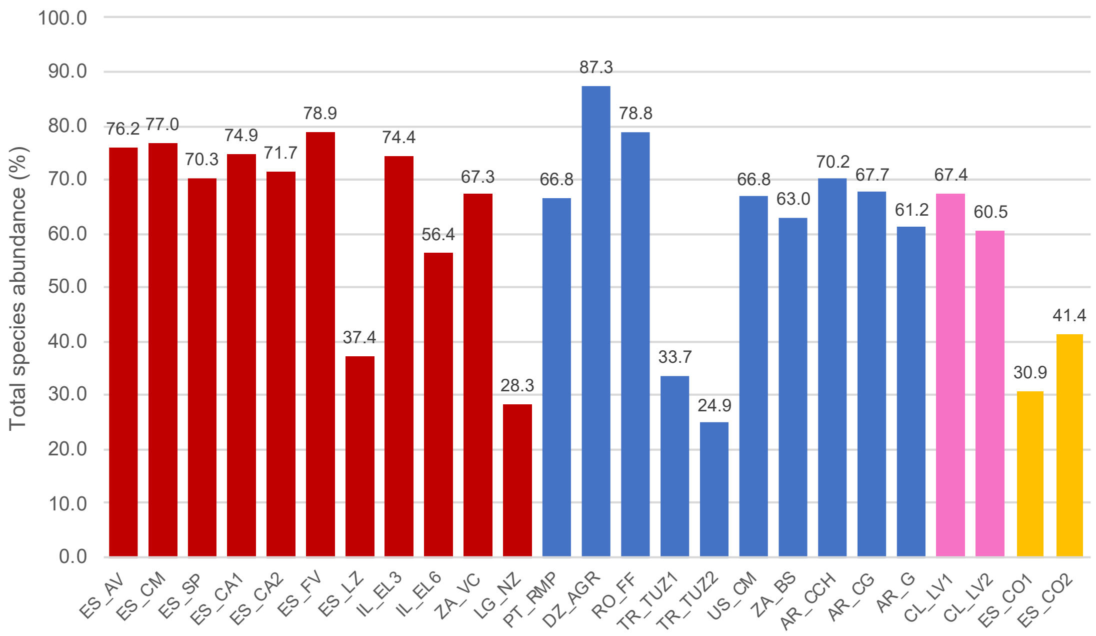

**Figure S9.** Similarity among (A) coastal and (B) inland samples based on Jaccard distance values compared to different geographic distances ranges. Jaccard distance index was evaluated based on presence or absence of the recovered species based on the MAG approach. The significant correlation between Jaccard dissimilarity and geographic distance was evaluated using a Procrustes test (correlation and p-value).

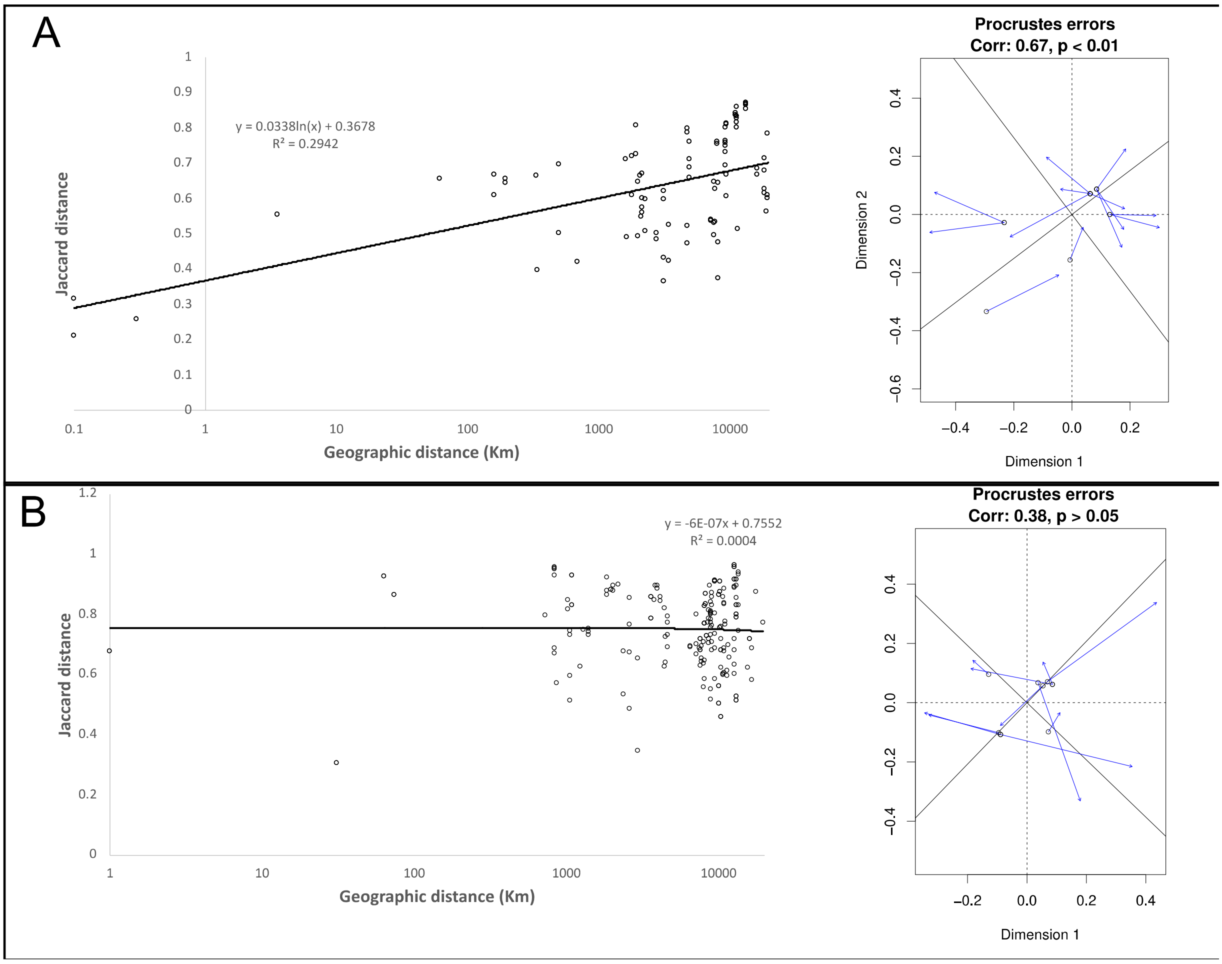

**Figure S10.** Similarity among all coastal salterns based on Jaccard distance in relation to pairwise geographic distance (kilometers): Dark red represents all pairwise comparisons of samples within a 0–400 km range, while standard red extends the analysis to include additional samples up to 5,000 km, and light red further extends the analysis to distances of 20,000 km.

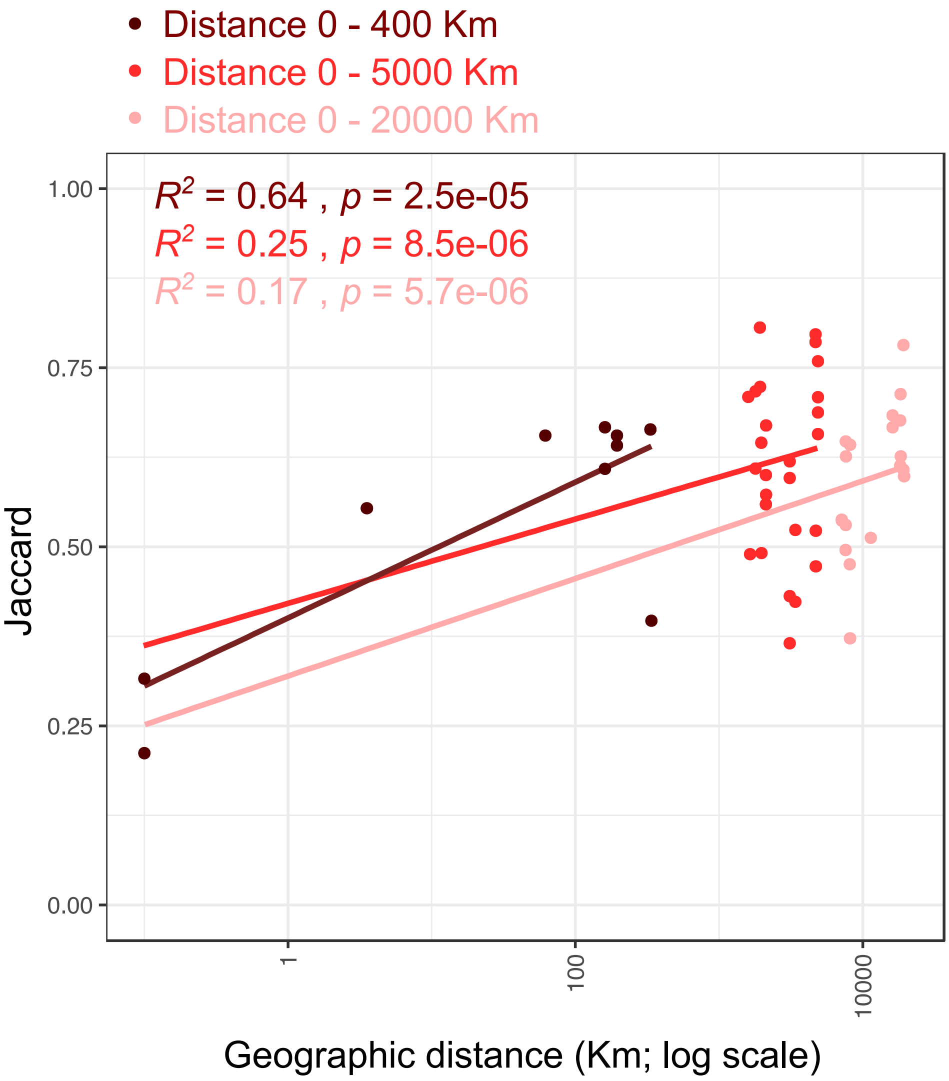

**Figure S11.** Similarity among inland hypersaline lakes based on Jaccard distance in relation to pairwise geographic distance (kilometers): Dark blue represents all pairwise comparisons of samples within a 0–1000 km range, while standard blue extends the analysis to include additional samples up to 5,000 km, and light blue further extends the analysis to distances of 20,000 km.

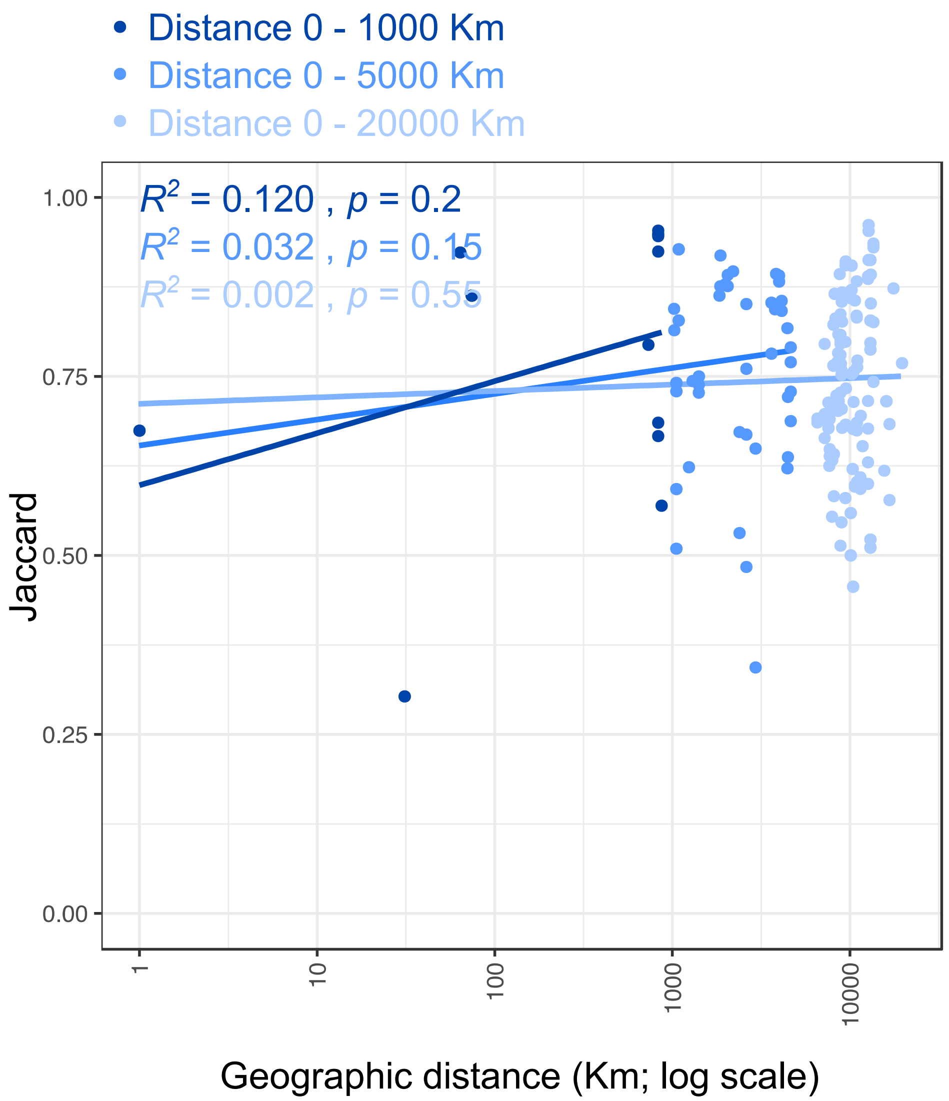

**Figure S12. Origin of MAGs from 25 species recovered across four or more metagenomic samples.** MAGs recovered from coastal solar salters are highlighted in read, while those from inland hypersaline lakes are shown in blue.

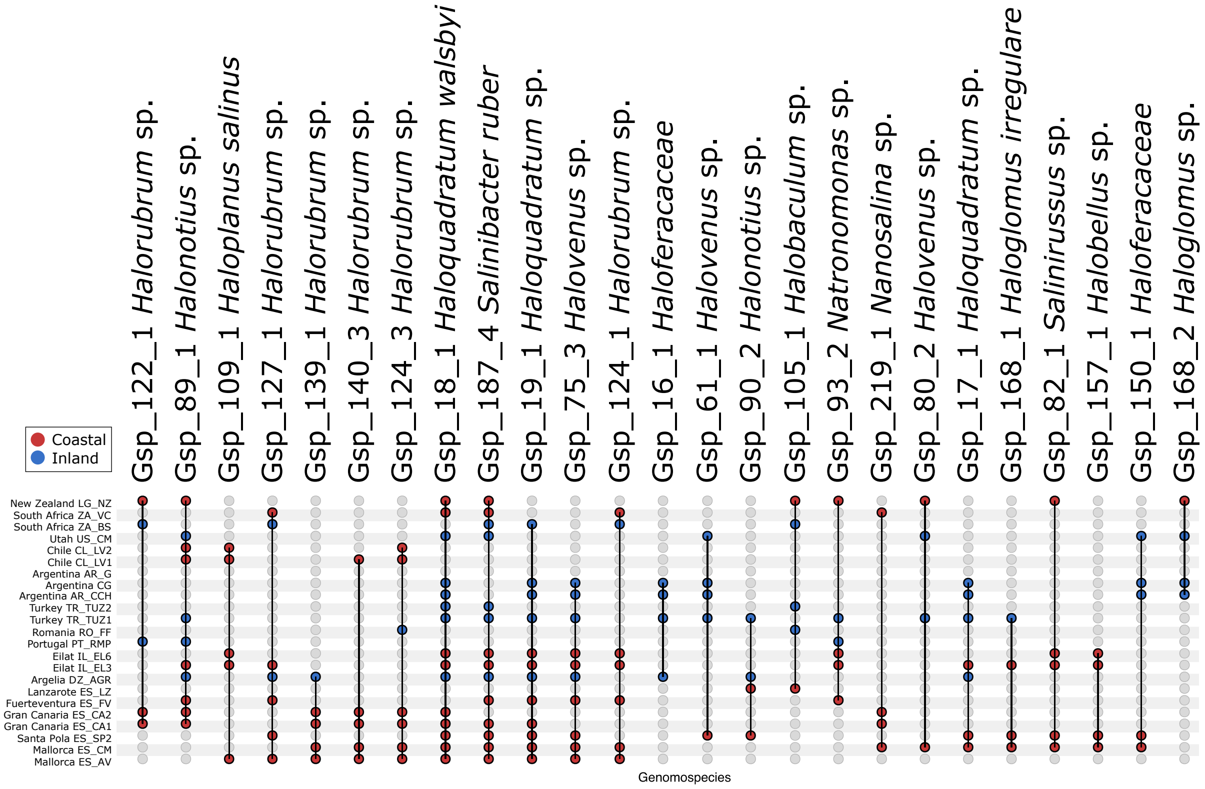

**Figure S13.** Intra-species average nucleotide identity (ANI, y-axes) compared to geographic distances (x-axis). (A) Species showing an ANI decrease with increasing geographic distance. (B) Species showing the decrease only in coastal salterns, but with no correlation among inland sites. (C) Highly conserved species with ANI values >99. (D) Species with no correlation between ANI and geographic distance.

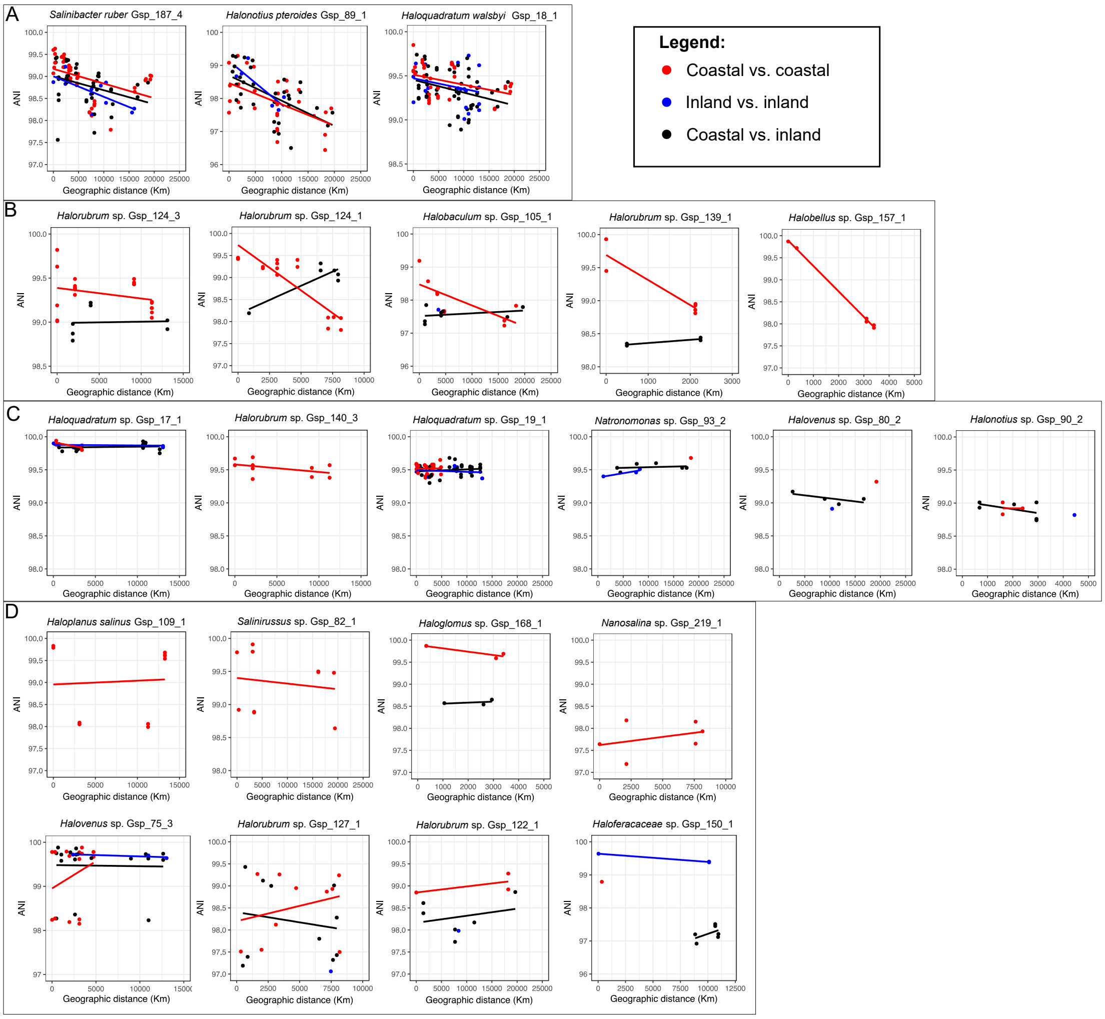

**Figure S14.** Intra-species average nucleotide identity (ANI, y-axes) compared to geographic distances (x-axis), considering just the intra-species core genome for ANI value calculations. (A) Species showing an ANI decrease with increasing geographic distance. (B) Species showing the decrease only in coastal salterns, but with no correlation among inland sites. (C) Highly conserved species with ANI values >99. (D) Species with no correlation between ANI and geographic distance.

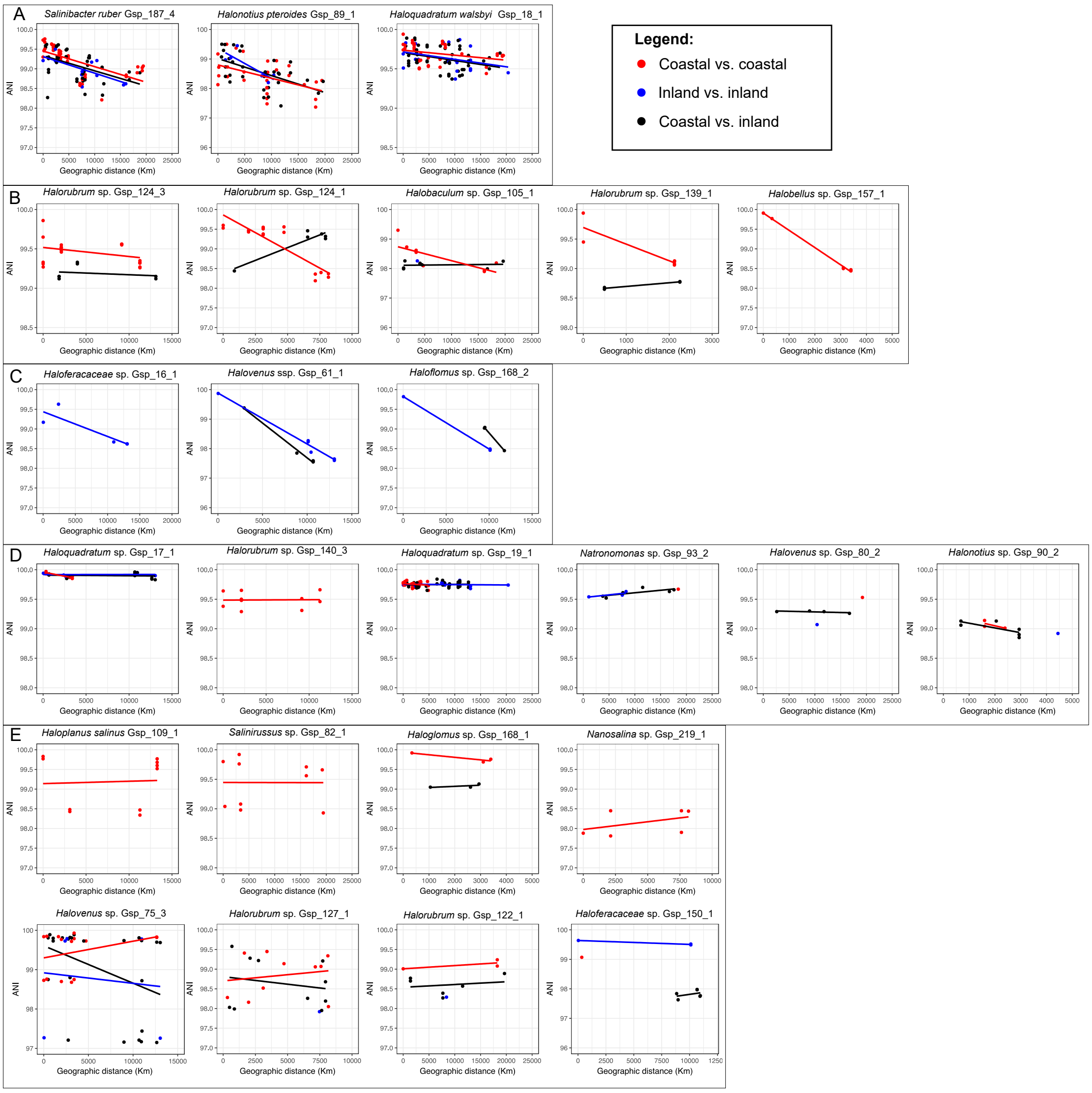

**Figure S15.** Intraspecies variation in core genes similarity across geographic distances, considering only MAGs from coastal sites for each species. To assess the impact of geographic distance on genomic divergence within species, we compared the sequence similarity of shared core genes between MAGs recovered from different sampling sites. Pairwise comparisons were grouped by geographic distance into short-range (0–400 km; red), intermediate-range (401–5000 km; green), and long-range (5001–20,000 km; blue) categories. Each plot displays the average distribution of core gene similarity values for all pairwise comparisons within each distance group. Each line represents the distribution of core-gene similarity values binned into 0.2% intervals. The four panels are organized according to Figure 5: (A) species showing an ANI decrease with increasing geographic distance; (B) species showing the decrease only in coastal salterns, but with no correlation among inland sites; (C) highly conserved species with ANI values >99; (D) species with no correlation between ANI and geographic distance.

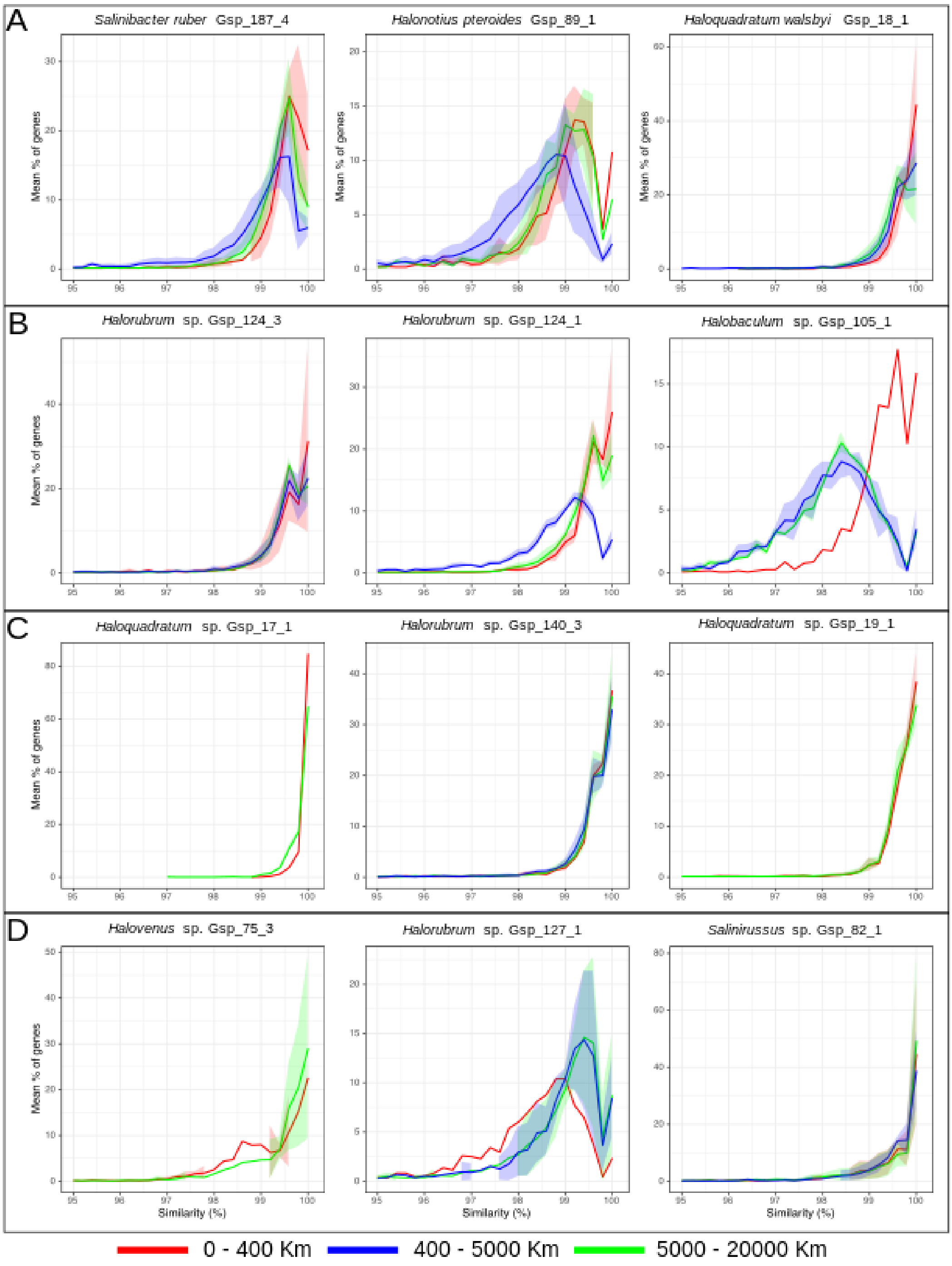

**Figure S16.** Relationship between pangenome-level similarity and geographic distance among MAGs of the same species. Relationship between geographic distance (x-axis) and gene-level similarity (y-axis, calculated as 1 – Bray-Curtis dissimilarity) for each species based on normalized gene abundance profiles (TAD80 values). Each point represents a pairwise comparison between MAGs, color-coded by sample origin: red for coastal–coastal comparisons, blue for inland–inland, and black for coastal–inland pairs. Linear regression lines are shown for coastal and inland comparisons when supported by sufficient data (n > 2). $R^2$ and p-values from the regressions are included in each plot. The four panels are organized according to Figure 5: (A) species showing an ANI decrease with increasing geographic distance; (B) species showing the decrease only in coastal salterns, but with no correlation among inland sites; (C) highly conserved species with ANI values >99; (D) species with no correlation between ANI and geographic distance.

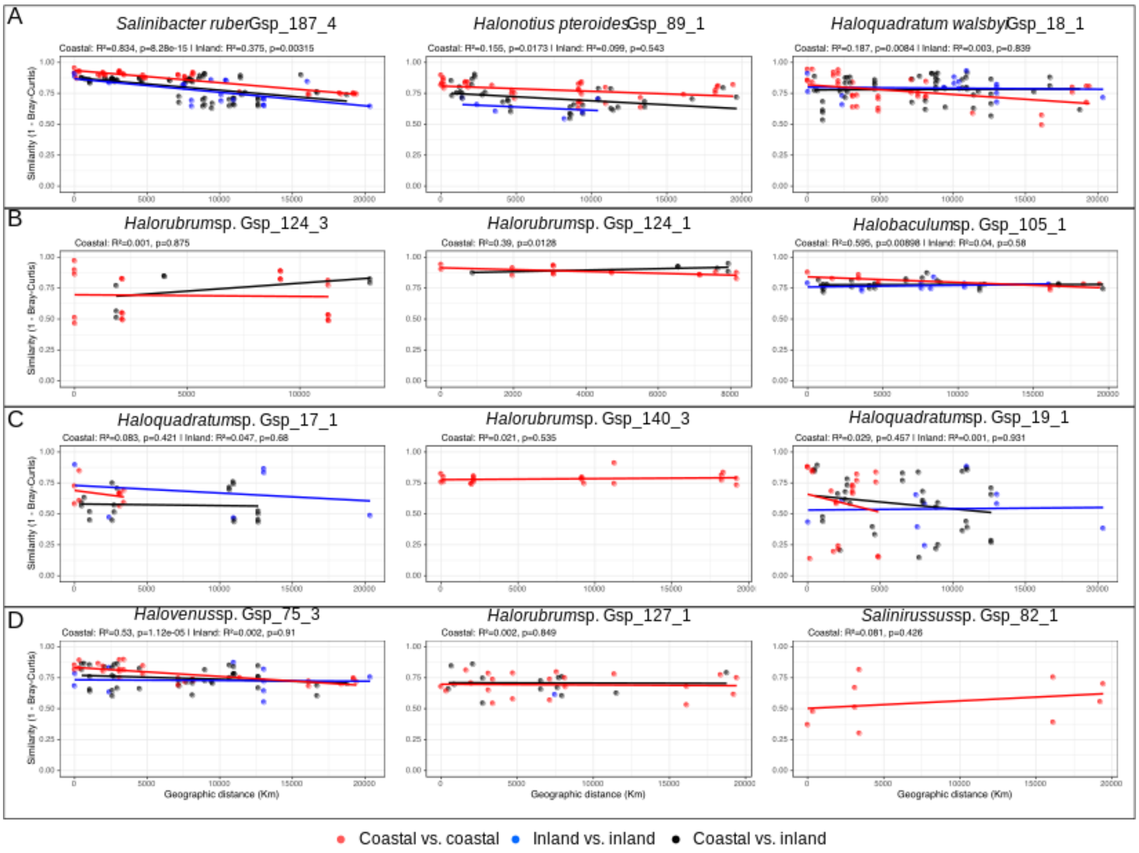

**Figure S17.** Relationship between pangenome-level similarity and geographic distance among MAGs of the same species. Relationship between geographic distance (x-axis) and gene-level similarity (y-axis, calculated as 1 – Jaccard dissimilarity), based on gene presence/absence profiles inferred from metagenomic read recruitment for each species. Each point represents a pairwise comparison between MAGs, color-coded by sample origin: red for coastal–coastal comparisons, blue for inland–inland, and black for coastal–inland pairs. Linear regression lines are shown for coastal and inland comparisons when supported by sufficient data (n > 2). $R^2$ and p-values from the regressions are included in each plot. The four panels are organized according to Figure 5: (A) species showing an ANI decrease with increasing geographic distance; (B) species showing the decrease only in coastal salterns, but with no correlation among inland sites; (C) highly conserved species with ANI values >99; (D) species with no correlation between ANI and geographic distance.

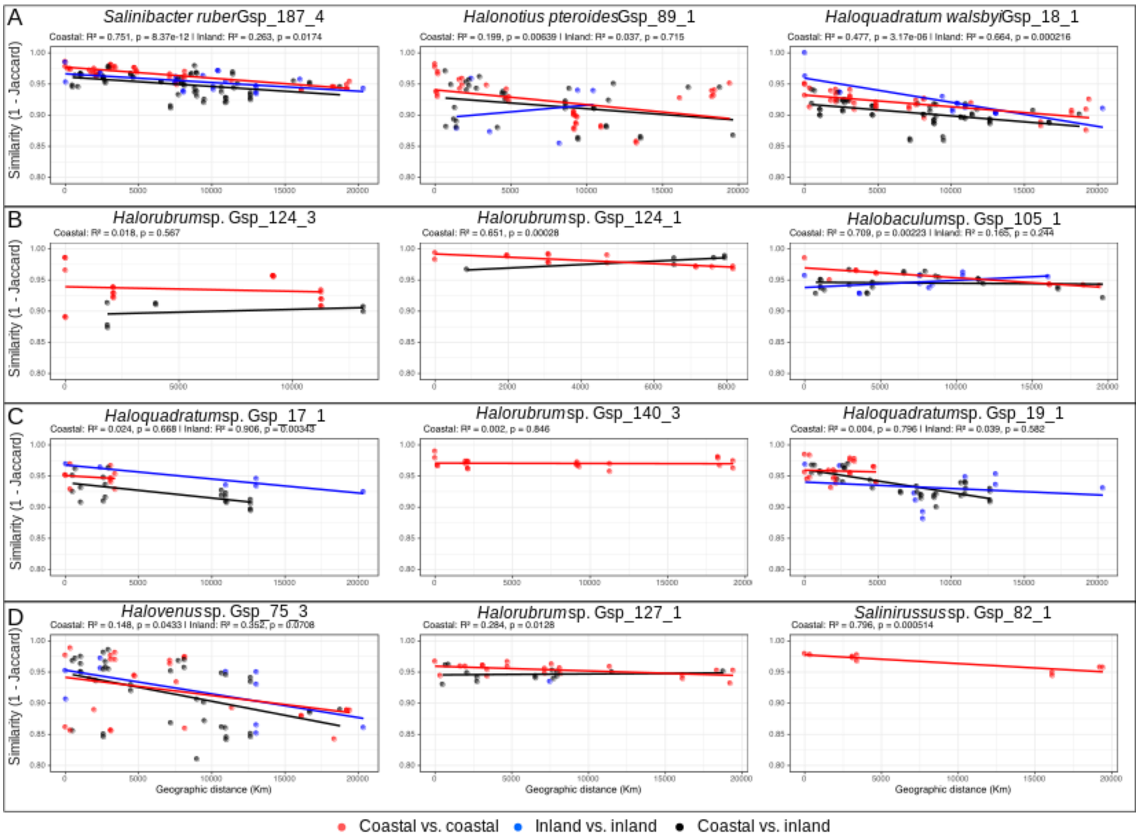

**Figure S18.** Functional categorization of genes with abundance patterns significantly correlated with geographic distance. For each species, pangenome gene abundance across samples was modeled against geographic distance using linear regression on all pairwise comparisons. Genes exhibiting significant positive or negative slopes ($p < 0.05$) were classified as showing increasing or decreasing abundance with distance, respectively. Barplots summarize the number of genes with significant trends across functional COG categories. Black bars represent genes whose abundance increased with distance, while grey bars indicate those that decreased. Genes without COG annotation are grouped under "Unc." (unclassified). The four panels are organized according to Figure 5: (A) species showing an ANI decrease with increasing geographic distance; (B) species showing the decrease only in coastal salterns, but with no correlation among inland sites; (C) highly conserved species with ANI values >99; (D) species with no correlation between ANI and geographic distance.

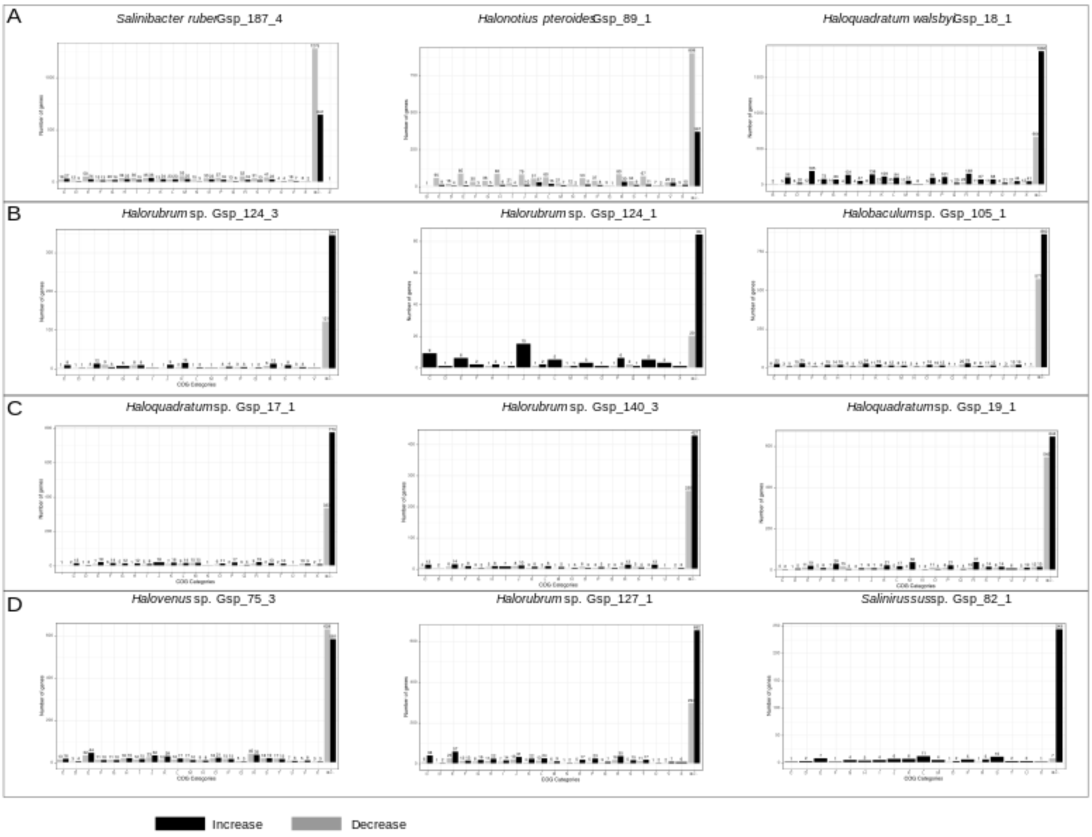

**Figure S19.** Functional enrichment of differentially abundant genes between coastal and inland samples. For each species, gene abundance profiles were compared between coastal and inland samples using the Wilcoxon rank-sum test. Genes with significant differences in abundance ($p < 0.05$) were classified as enriched in either coastal or inland environments based on the direction of the log2 fold change. These differentially abundant genes were annotated with COG functional categories, and their distribution was summarized to identify overrepresented functions in each group. Plots display the number of enriched genes per COG category for coastal (red) and inland (blue) samples. Categories significantly enriched in one group over the other (Fisher's exact test, $p < 0.05$) are marked with asterisks. Genes without COG annotation were grouped under "Unc." (unclassified). The four panels are organized according to Figure 5: (A) species showing an ANI decrease with increasing geographic distance; (B) species showing the decrease only in coastal salterns, but with no correlation among inland sites; (C) highly conserved species with ANI values >99; (D) species with no correlation between ANI and geographic distance.

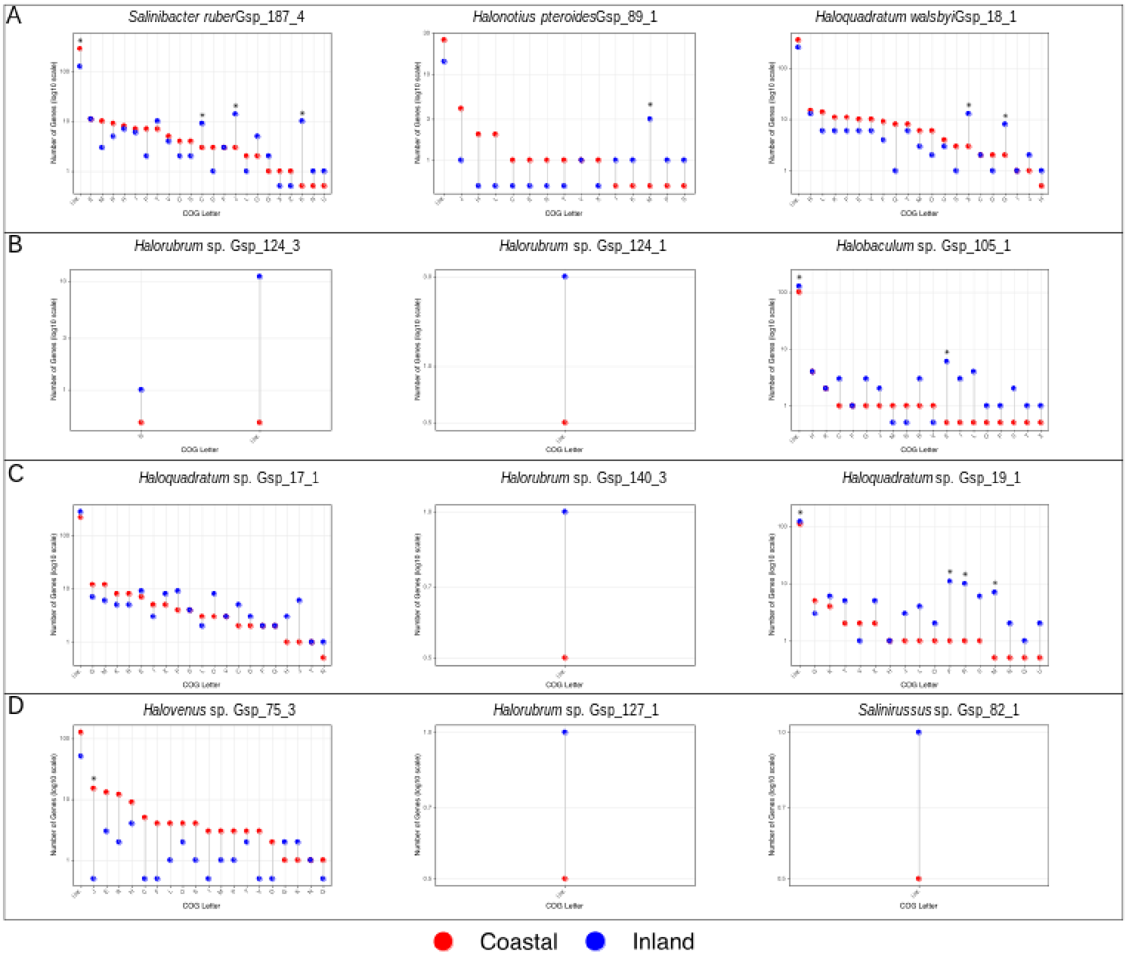

**Figure S20.** Sampling of hypersaline brines from Coco pools (Mallorca Island, Spain) and Lo Valdivia solar salterns (Chile). (A) Geographic location of Coco pools on the southern rocky coast of Mallorca Island; (B) laboratory-processed brine samples in bottles; (C, D, E) field sampling of brine from Coco 1 and Coco 2 pools; (F) brine pellets obtained after centrifugation, used for DNA extraction and metagenomic sequencing. (G, H) Lo Valdivia salterns during the winter season, when ponds are flooded by seawater intrusions from the nearby river during high tide; (I, J) Lo Valdivia salterns in summer, showing the formation of crystallizer ponds by seawater evaporation and concentration of brines.

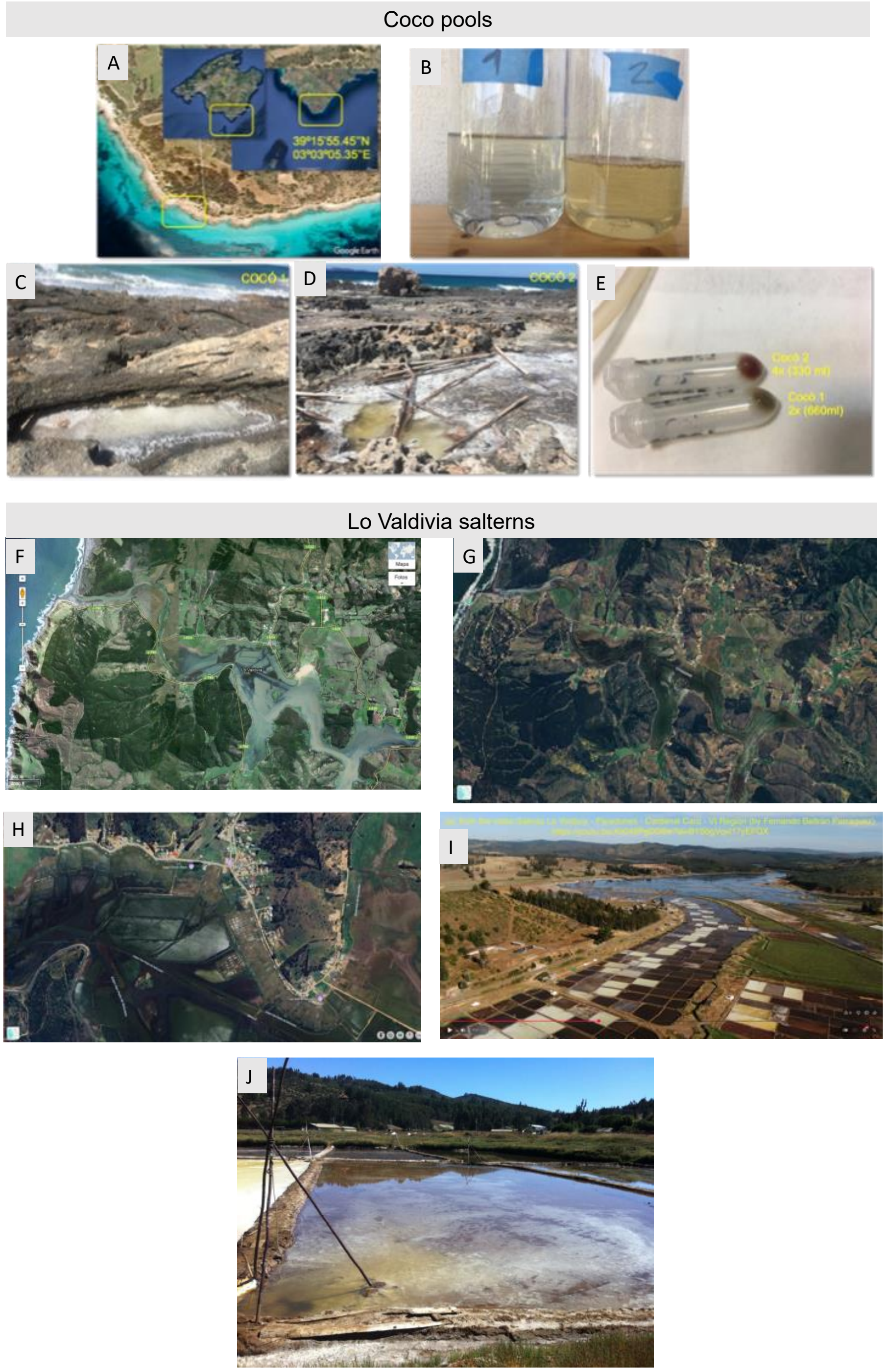

**Figure S21.** Similarity among all metagenomes determined in this study, including those samples from Coco coastal pools. NMDS analysis of Jaccard-based distances using MAGs diversity patterns.

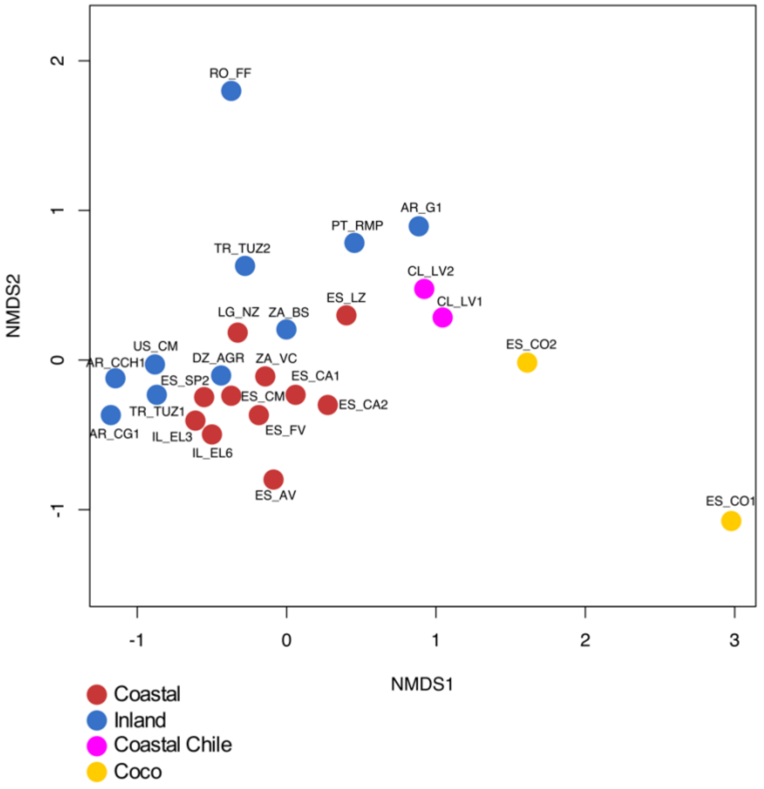

**Figure S22.** Complex upset graph showing the shared MAGs between the Coco samples and the other coastal and inland samples included in this study.

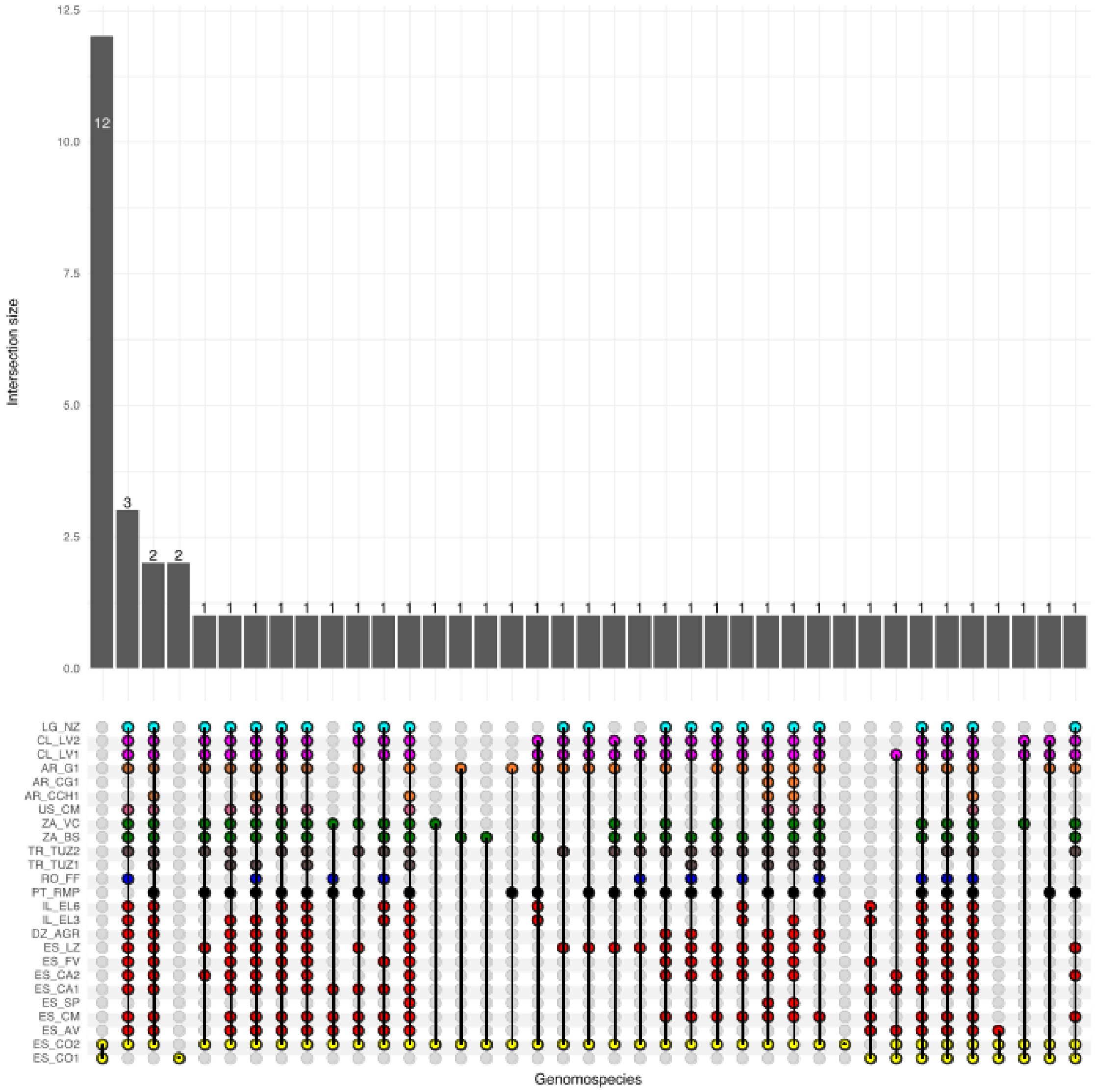

**Figure S23.** Number of MAGs shared between Coco with coastal, inland and Chilean (Lo Valdivia) metagenomes. (A) Barplot indicating the number of MAGs shared between coco and all the other samples. (B) Boxplot differentiated by coastal solar saltern, inland hypersaline lakes and Lo Valdivia samples. (C) Boxplot comparing the Mash distance dissimilarity values between the Coco samples to coastal, inland or the Chilean samples. (D) Boxplot comparing the Mash distance dissimilarity between the Chilean samples to coastal, inland or the Coco samples.

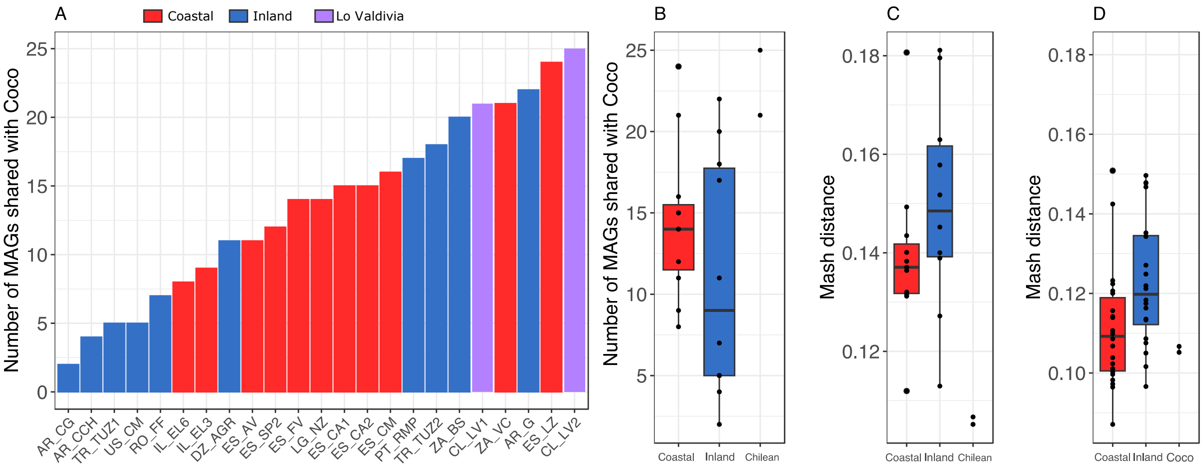
